# Broadly neutralizing anti-S2 antibodies protect against all three human betacoronaviruses that cause severe disease

**DOI:** 10.1101/2022.03.04.479488

**Authors:** Panpan Zhou, Ge Song, Wan-ting He, Nathan Beutler, Longping V. Tse, David R. Martinez, Alexandra Schäfer, Fabio Anzanello, Peter Yong, Linghang Peng, Katharina Dueker, Rami Musharrafieh, Sean Callaghan, Tazio Capozzola, Meng Yuan, Hejun Liu, Oliver Limbo, Mara Parren, Elijah Garcia, Stephen A. Rawlings, Davey M. Smith, David Nemazee, Joseph G. Jardine, Ian A. Wilson, Yana Safonova, Thomas F. Rogers, Ralph S. Baric, Lisa E. Gralinski, Dennis R. Burton, Raiees Andrabi

## Abstract

Pan-betacoronavirus neutralizing antibodies may hold the key to developing broadly protective vaccines against coronaviruses that cause severe disease, for anticipating novel pandemic-causing viruses, and to respond more effectively to SARS-CoV-2 variants. The emergence of the Omicron variant of SARS-CoV-2 has illustrated the limitations of solely targeting the receptor binding domain (RBD) of the envelope Spike (S)-protein. Here, we isolated a large panel of broadly neutralizing antibodies (bnAbs) from SARS-CoV-2 recovered-vaccinated donors that target a conserved S2 region in the fusion machinery on betacoronavirus spikes. Select bnAbs show broad *in vivo* protection against all three pathogenic betacoronaviruses, SARS-CoV-1, SARS-CoV-2 and MERS-CoV, that have spilled over into humans in the past 20 years to cause severe disease. The bnAbs provide new opportunities for antibody-based interventions and key insights for developing pan-betacoronavirus vaccines.

## Main

The initial successes of SARS-CoV-2 vaccines were due in part to the relative ease of inducing protective neutralizing antibodies (nAbs) to immunodominant epitopes on the receptor binding domain (RBD) of the S1 subunit of the spike protein (*1-5*). The most potent nAbs target epitopes overlapping the ACE2 receptor binding site (RBS) (RBS-A/class 1; RBS-B,C,D/ class 2 antibodies) and require little affinity maturation to neutralize at very low (single digit ng/ml) concentrations (*3, 6-12*). However, mutations, particularly in and around the RBS, readily generate viral variants that are resistant to neutralization by commonly induced classes of antibodies (*13, 14*) without significantly negatively impacting viral fitness, and have led to several Variants of Concern (VOCs) (*14-25*). For example, the K417N/T and E484K mutations in the Beta and Gamma VOCs leads to neutralization escape from the vast majority of RBS-A/class 1 and RBS-B, C, D/class 2 nAbs (*12-14*). Other sarbecoviruses using ACE2 as receptor, such as SARS-CoV-1, and betacoronaviruses using receptors other than ACE2, such as MERS-CoV, show even more sequence divergence in the RBS region (*26-28*). The emergence of SARS-CoV-2 VOCs, together with a desire to have the capability to respond to novel coronaviruses with pandemic potential, has focused effort on vaccines and antibodies that target the most conserved regions of the spike protein (*29-31*). The “lower” more conserved faces of the RBD have been investigated and are targeted by many nAbs with greater breadth of neutralization against SARS-CoV-2 variants and diverse sarbecoviruses than for example RBS-A/class 1 or RBS-B,C,D/class 2 nAbs (*32-41*). However, the Omicron variant has demonstrated escape also from some nAbs targeting these more conserved regions of the RBD (*19, 25*). An alternative relatively conserved target on the coronavirus spike is the S2 region, which does harbor neutralizing epitopes (*42*) and therefore is of interest for attempts to generate SARS-CoV-2 vaccines effective against VOCs and, more ambitiously, pan-betacoronavirus vaccines (*29-31*).

We recently isolated a nAb, CC40.8, from a COVID-19 convalescent donor that neutralized sarbecoviruses from clades 1a and 1b and SARS-CoV-2 VOCs (*43, 44*). We demonstrated antibody protection against SARS-CoV-2 challenge in human ACE2 mouse and hamster models (*44*). We further mapped the epitope of CC40.8 to the conserved spike S2 stem-helix region that forms part of the spike fusion machinery (*44*). Several more bnAbs targeting this region have been isolated from humans and from vaccinated animals (*45-51*). These bnAbs are a good starting point and highlight the opportunities that conserved bnAb S2 epitopes may offer for broad betacoronavirus vaccine targeting. However, a large panel of stem-helix bnAbs will be needed to define the common molecular features of antibodies targeting this site, which may facilitate the development of rational vaccine strategies that can induce such bnAbs by vaccination (*52-56*). Such a panel would also provide more options for antibody-based prophylaxis and therapeutic strategies (*57*).

## Results

### Donors for isolation of β-CoV spike stem-helix bnAbs

To identify suitable donors for the isolation of a panel of β-CoV spike stem-helix bnAbs, we screened immune sera from human donors for cross-reactive binding to 25-mer spike stem-helix region peptides, which we previously identified as a target for bnAbs (*43, 44*). We tested sera from three different groups of donors: i) COVID-19 recovered donors (n = 15); ii) spike mRNA-vaccinated (2X) donors (n = 10) and iii) COVID-19-recovered then spike-vaccinated (1X) donors (n = 15) (Fig. 1a). Whereas weak or no binding was observed for COVID-19 recovered or vaccinee sera to human β-CoV spike stem-helix peptides, sera from 80% (12/15) of recovered-vaccinated donors exhibited strong cross-reactive binding to the peptides (Fig. 1a). We noted a strong correlation between binding of recovered-vaccinated sera to SARS-CoV-2 stem-helix peptide with binding to other human β-CoV stem-helix peptides suggesting targeting of common cross-reactive epitopes (Fig. 1b). Accordingly, we sought to isolate β-CoV stem-helix directed bnAbs from 10 SARS-CoV-2 recovered-vaccinated donors that exhibited cross-reactive binding to this spike region.

**Figure. 1.**
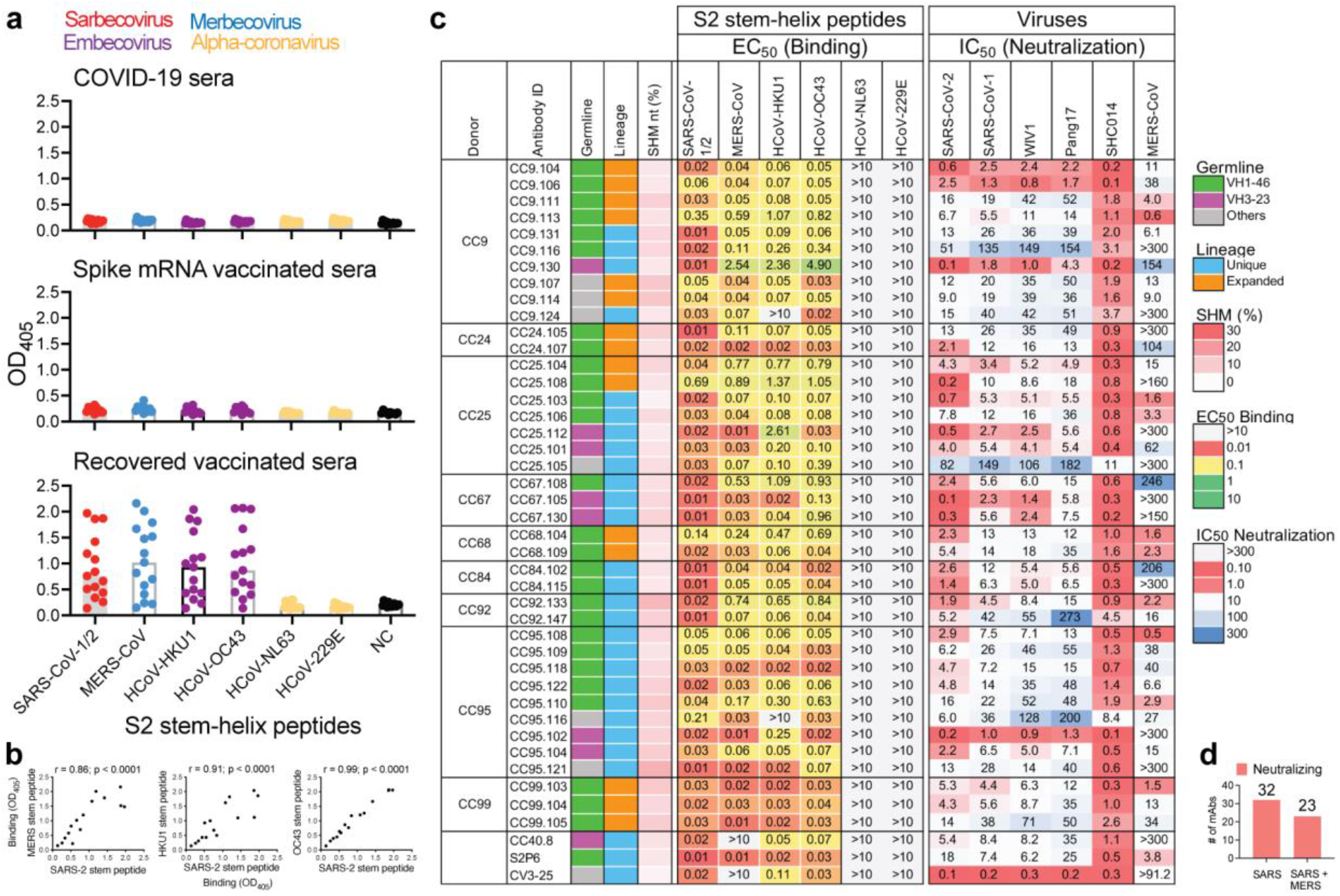
Binding and neutralization properties of S2 stem-helix mAbs. **a.** Dot plots showing ELISA binding (OD_405_) reactivity of immune sera from COVID-19 convalescent donors (n = 15), spike mRNA-vaccinated donors (n = 10) and SARS-CoV-2 recovered-vaccinated donors (n = 15) to 25-mer peptides corresponding to spike S2 stem-helix regions of human β-(sarbecoviruses: SARS-CoV-1 or 2; merbecovirus: MERS-CoV; embecoviruses: HCoV-HKU1, HCoV-OC43) and α-(HCoV-NL63 and HCoV-229E) coronaviruses. 12 out of 15 (80%) SARS-CoV-2 recovered-vaccinated donor sera show cross-reactive binding to β-CoV spike stem-helix peptides. **b.** Correlation between binding of infected-vaccinated sera to SARS-CoV-2 stem-helix peptide and the other β-CoV (MERS-CoV, HCoV-HKU1 and HCoV-OC43) stem-helix peptides. Responses for binding to two stem-helix peptides were compared by nonparametric Spearman correlation two-tailed test with 95% confidence interval and the Spearman correlation coefficient (r) and the p-value are indicated. **c.** A total of 40 S2 stem-helix mAbs were isolated from 9 SARS-CoV-2 recovered-vaccinated donors (CC9, CC24, CC25, CC67, CC68, CC84, CC92, CC95 and CC99). MAbs were isolated by single B cell sorting using SARS-CoV-2 and MERS-CoV S-proteins as baits. Heatmap showing IGVH germline gene usage (colored: VH1-46 (green), VH3-23 (plum) and other V-genes (grey)), lineage information (unique (sky) and expanded (tangerine) lineages) and V-gene nucleotide somatic hypermutations (SHMs). EC_50_ ELISA binding titers of mAbs with β- and α-HCoV spike S2 stem-helix region peptides. MAbs showed binding to β-but not α-HCoV derived stem-helix peptides. IC_50_ neutralization of mAbs against pseudoviruses of clade1a (SARS-CoV-2 and Pang17), clade 1b (SARS-CoV-1, WIV1, SHC014) sarbecoviruses and MERS-CoV. Spike S2 stem-helix bnAbs, CC40.8, S2P6 and CV3-25 were used as controls for binding and neutralization assays. **d.** 32 of 40 stem-helix bnAbs were unique clones that neutralized all ACE2-utilizing sarbecoviruses and 23 out of 32 unique mAb neutralized MERS-CoV, in addition to sarbecoviruses.

### Isolation of a large panel of β-CoV spike stem-helix mAbs

Using SARS-CoV-2 and MERS-CoV S-proteins as baits, we sorted antigen-specific single B cells to isolate 40 stem-helix mAbs from 10 COVID-19 convalescent donors who had been recently vaccinated with the Pfizer/BioNTech BNT162b2 (n = 4: CC9, CC92, CC95 and CC99), Johnson & Johnson Ad26.CoV2.S (n = 1: CC67), or Moderna mRNA-1273 (n = 5: CC24, CC25, CC26, CC67, CC84) vaccines (Fig. 1c, d, Supplementary Fig. 1) (*2, 58, 59*). Briefly, using SARS-CoV-2 and MERS-CoV S-proteins, we sorted CD19^+^CD20^+^IgG^+^IgM^−^ B cells positive for both probes from the peripheral blood mononuclear cells (PBMCs) of these donors. Flow cytometry profiling revealed up to 36% (range = 6 – 36%, median = 15%) SARS-CoV-2 S-protein-specific B cells, of which a sizable fraction was cross-reactive with the MERS-CoV S-protein (range = 0.04 – 0.28%, median = 0.16% total selected B cells) (Supplementary Fig. 1b). A total of 358 SARS-CoV-2: MERS-CoV S-protein-specific double positive single B cells were recovered from the 10 donors, of which the heavy (HC)-light (LC) chain pairs were recovered from 247 single B cells (69%) from 9 donors and expressed as IgGs (Supplementary Fig. 1c). Expi293F cell-expressed IgG supernatants of 247 mAbs were screened for dual binding to SARS-CoV-2 and MERS-CoV stem-helix peptides and 16% (40/247) exhibited cross-reactive binding (Supplementary Fig. 1c). Dual binding was confirmed for the corresponding purified IgGs. Except for two mAbs that failed to bind HCoV-HKU1 stem-helix peptide, all mAbs exhibited cross-reactive binding to stem-helix peptides of endemic β-HCoV (HCoV-HKU1 and HCoV-OC43) but not α-HCoV (HCoV-NL63 and HCoV-229E) (Fig. 1c). We also tested binding of mAbs to soluble HCoV S-proteins and cell surface expressed spikes and observed consistent binding to SARS-CoV-2/1 and MERS-CoV spikes but reduced binding to endemic β-HCoV spikes (HCoV-HKU1 and HCoV-OC43), especially in the soluble S-protein format (Supplementary Fig. 2). Overall, we isolated 40 stem-helix mAbs, of which 32 were encoded by unique immunoglobulin germline gene combinations and 7 were expanded lineages with 2 or more clonal members (Fig. 1c, Supplementary Fig. 2).

### Spike stem-helix mAbs exhibit broad neutralization against β-CoVs

We next examined neutralization of stem-helix mAbs against clade 1a (SARS-CoV-1, WIV1 and SHC014) and clade 1b (SARS-CoV-2 and Pang17) ACE2-utilizing sarbecoviruses (*26, 27*) and MERS-CoV (*28*). Consistent with conservation of the stem-helix bnAb epitope region across sarbecoviruses, all the 32 mAb lineages neutralized all the 5 sarbecoviruses tested with widely varying degrees of neutralization potency (Fig. 1c, d). The bnAbs neutralized clade 1a SHC014 and clade 1b SARS-CoV-2 relatively more potently compared to the other sarbecoviruses, but some bnAbs neutralized all viruses in the lower μg/mL IC_50_ neutralization titer range (0.1 to 3 μg/ml). Of 32 unique stem-helix bnAb lineages, 23 (72%) bnAbs neutralized MERS-CoV (Fig. 1c, d). Neutralization potency against MERS-CoV was lower compared to the sarbecoviruses but many bnAb members were consistently effective. We tested neutralization of SARS-CoV-2 VOCs (B.1.1.7 (Alpha), B.1.351 (Beta), P.1 (Gamma), B.1.617.2 (Delta) and B.1.1.529 (Omicron) by select bnAbs (Fig. 2a). Consistent with the conservation of the stem-helix region in SARS-CoV-2 VOCs, these bnAbs were consistently effective against the VOCs tested (Fig. 2a). Of note, a fraction of stem-helix bnAbs showed some degree of polyreactivity or autoreactivity in HEp2 cell or polyspecificity reagent (PSR) assays (*7*) but the majority were negative (Supplementary Figs. 2 & 4). Overall, we have identified multiple stem-helix bnAbs that exhibit broad neutralizing activity against phylogenetically diverse β-HCoVs.

**Figure. 2.**
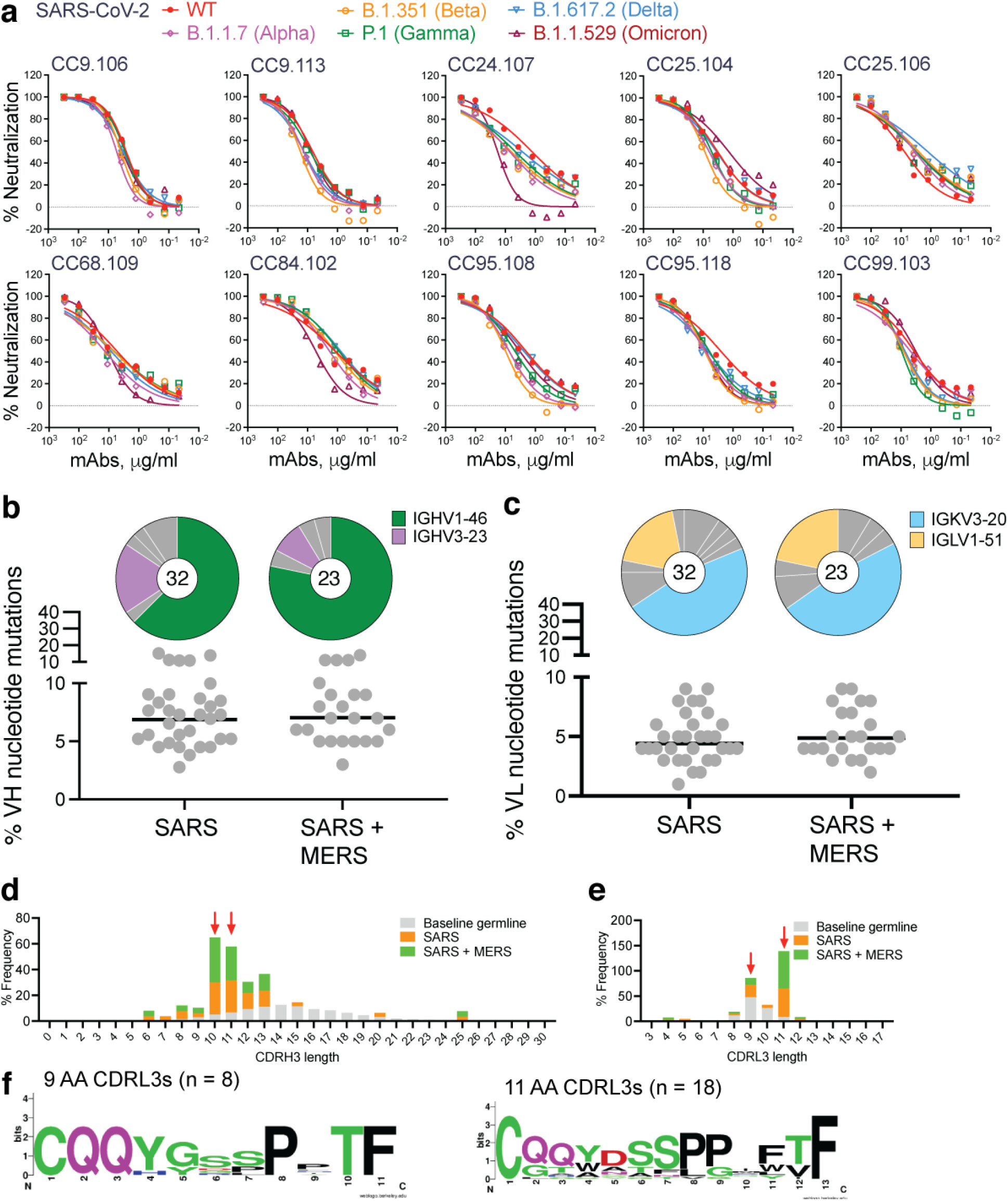
Neutralization of SARS-CoV-2 VOCs, and immunogenetic properties of S2 β-CoV spike stem-helix bnAbs. **a.** Neutralization of 10 select S2 stem-helix bnAbs against SARS-CoV-2 (WT) and five major SARS-CoV-2 variants of concern [B.1.1.7 (Alpha), B.1.351 (Beta), P.1 (Gamma), B.1.617.2 (Delta), and B.1.1.529 (Omicron)]. **b-c.** Pie plots showing IGHV and IGKV/IGLV gene usage distribution of isolated stem-helix mAbs. Enriched heavy (IGHV1-46 (green) and IGHV3-23 (plum)) **(b)** and light (IGKV3-20 (sky) and IGLV1-51 (cantaloupe)) **(c)** gene families are colored. Dot plots showing % nucleotide mutations (SHMs) in the heavy (VH) or light (VL) chains of isolated stem-helix mAbs. The mAbs are grouped by neutralization against sarbecoviruses or sarbecoviruses + MERS-CoV. **d-e.** CDRH3 **(d)** or CDRL3 **(e)** length distributions of isolated mAbs across sarbecovirus broadly neutralizing and sarbecovirus + MERS-CoV broadly neutralizing mAb groups compared to human baseline germline reference. MAbs with 10- and 11-amino acid-CDRH3s or mAbs with 9- and 11-amino CDRL3s, enriched in S2 stem-helix bnAbs compared to baseline germline reference, are indicated by red arrows. **f.** Sequence conservation logos of 9 (n = 8) and 11 (n = 18) amino acid long CDRL3-bearing stem-helix bnAbs show enrichment of certain J-gene encoded residues.

### Immunogenetics of stem-helix bnAbs and vaccine targeting

Immunogenetic analysis of stem-helix antibody sequences showed strong enrichment of IGHV1-46 (63%) and IGHV3-23 (22%) germline gene families as compared to human baseline germline frequencies (Figs. 1c, 2b, Supplementary Fig. 3) (*60, 61*). Of note, previously isolated stem-helix human bnAbs, S2P6 and CC40.8, are IGHV1-46 and IGHV3-23 germline encoded, respectively (*44, 45*). The IGHV1-46 germline gene was slightly more enriched (78%) in stem-helix bnAbs that exhibited MERS-CoV neutralization in addition to sarbecoviruses, suggesting a potential role for this VH-germline gene for broader reactivity against diverse β-HCoV spikes. Interestingly, at least one IGHV1-46-encoded stem-helix bnAb was isolated from each of the 9 donors and may represent a public clonotype for this bnAb site. For light chain gene usage, we noted a strong enrichment of IGKV3-20 (47%) and to some degree IGLV1-51 (16%) germline gene families as compared to human baseline germline frequencies (Figs. 1c, 2c, Supplementary Fig. 3) (*62*). The mAbs possessed modest levels of V-gene nucleotide somatic hypermutation (SHM): for VH, median = 7.3% and for VL, median = 4.5% (fig. S2).

We examined the CDRH3 loop lengths in the isolated stem-helix bnAbs and observed a strong enrichment for 10- and 11-residue long CDRH3s compared to the human baseline reference database (Fig. 2d, Supplementary Fig. 3) (*60, 61*). No apparent enrichment in germline D-genes was observed but IGHJ4, the most common germline J-gene utilized in humans, was slightly enriched (72%) in stem-helix bnAbs compared to a reference germline database (Supplementary Fig. 3) (*60, 61*). We also examined the CDRL3 loop lengths in the stem-helix bnAbs and observed strong enrichment for 9- and 11-residue CDRL3s (Fig. 2e, Supplementary Figs. 2 and 3). These CDRL3 loops possess germline JL-gene-encoded motifs (Fig. 2f, Supplementary Fig. 2), which may be important for epitope recognition. Overall, we observed a strong enrichment of IGHV and IGLV germline gene features in β-HCoV spike stem-helix bnAbs. Therefore, rational vaccine strategies may exploit these germline gene features to generate a protective B cell response (*53, 54, 63*).

To examine the potential contribution of antibody SHMs to SARS-CoV-2 neutralization efficiency and cross-neutralization with MERS-CoV, we tested the binding of select mAbs (based on a broad range of neutralization potency) to SARS-CoV-2 or MERS-CoV monomeric stem-helix peptides and to their S-proteins by BLI (Supplementary Fig. 5). The mAbs bind SARS-CoV-2 and MERS-CoV-2 stem-helix peptides with nanomolar (nM) and higher K_D_ affinity (Supplementary Fig. 5a) and were generally higher for SARS-CoV-2 compared to MERS-CoV stem-helix peptide. We found no association of heavy or light chain SHMs with binding to SARS-CoV-2 or MERS-CoV-2 stem-helix peptides or with neutralization of the corresponding viruses (Supplementary Fig. 5b). We however observed a strong association of binding affinity to stem-helix peptides and neutralization (Supplementary Fig. 5c).

To further investigate the role of SHM in binding and neutralization, we generated inferred germline (iGL) versions of stem-nAbs by reverting their heavy and light chain V, D and J regions to the corresponding germlines (inferred germlines, iGLs) as described previously (*64*) and assessed both binding and neutralization. The BLI binding responses and the K_D_ values of the bnAb iGLs with SARS-CoV-2 and MERS-CoV stem-helix peptides were substantially reduced compared to mature bnAbs but were still strong and in the lower nM and higher K_D_ affinity range (Fig. 3a, Supplementary Fig. 5a). We observed higher affinities or CDRH3 “RG” motif-bearing IGHV1-46-encoded and CDRL3 “WD” motif-bearing IGLV1-51-encoded bnAb iGLs for binding to SARS-CoV-2 or MERS-CoV stem-helix peptides (Fig. 3a, Supplementary Fig. 6). Binding of bnAbs and their iGLs to S-proteins were generally of higher affinity than to the corresponding peptides, possibly due to avidity effects (Supplementary Fig. 5a). The affinities of iGLs compared to mature bnAbs were notably less for S-proteins compared to the corresponding peptides, particularly for the MERS-CoV S-protein where many of the iGL Abs failed to bind substantially (Supplementary Fig. 5a). Overall, these results suggest a significant contribution from germline-encoded residues to epitope binding, in most cases consistent with enrichment of certain antibody germline gene features above (Figs. 1 and 2).

**Figure. 3.**
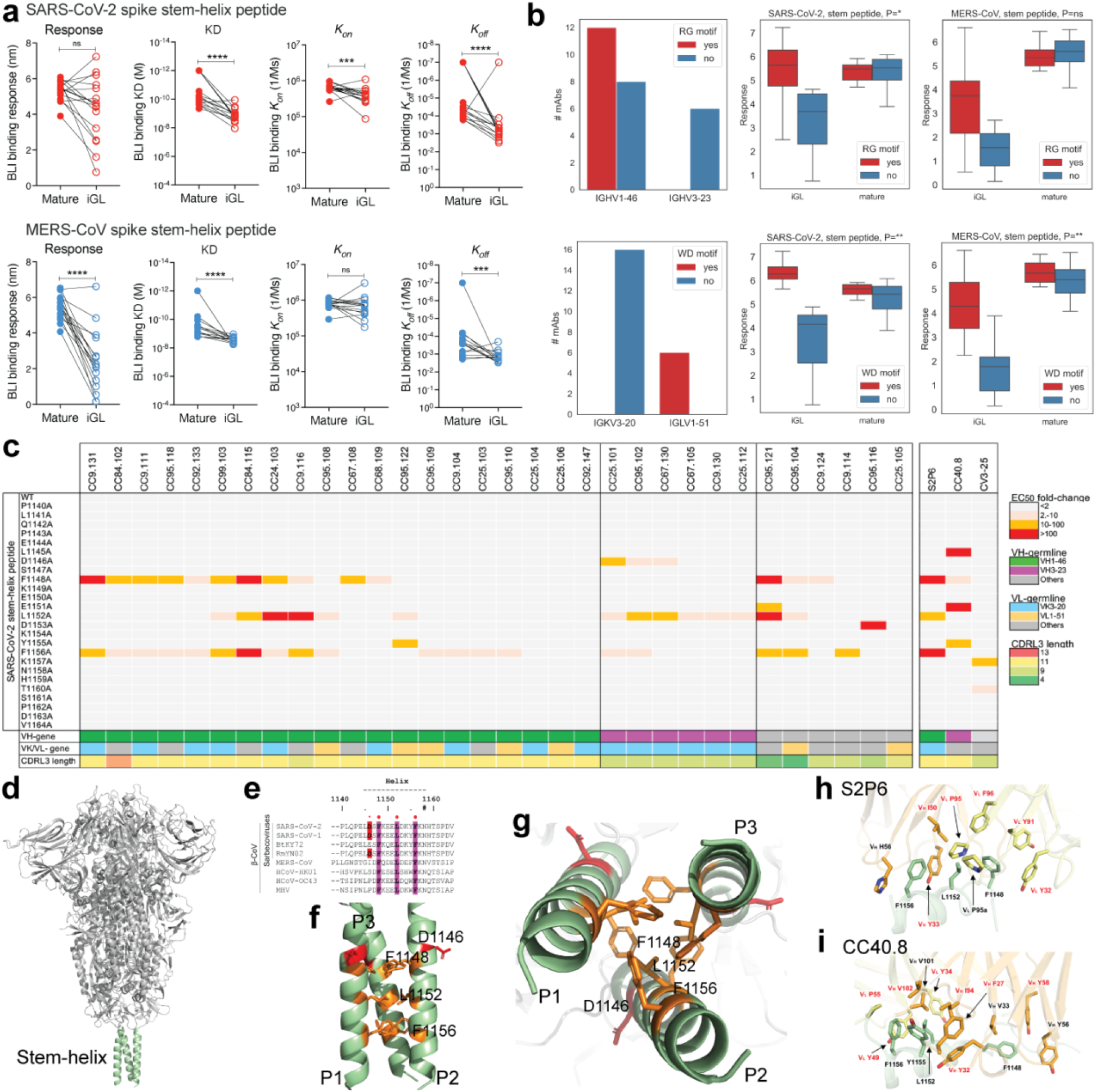
Binding kinetics and fine epitope specificities of S2 β-CoV spike stem-helix bnAbs. **a.** BLI binding kinetics of S2 stem-helix mature bnAbs and their inferred germline (iGL) versions to monomeric SARS-CoV-2 and MERS-CoV stem-helix peptides. Binding kinetics were obtained using a 1:1 binding kinetics fitting model on ForteBio Data Analysis software and maximum binding responses, dissociations constants (*K*_D_) and on-rate (*k_on_*) and off-rate constants (*k_off_*) for each antibody-protein interaction are compared. K_D_, k_on_ and k_off_ values were calculated only for antibody-antigen interactions where a maximum binding response of 0.2nm was obtained. The S2 stem-helix bnAb iGL Abs showed substantially reduced binding to stem-helix peptides compared to their corresponding mature versions. Statistical comparisons between two groups were performed using a Mann-Whitney two-tailed test, (***p < 0.001, ****p < 0.0001; ns- p >0.05). **b.** Association of S2 stem-helix peptide binding by S2 bnAbs and their iGLs with CDRH3 and CDRL3 motifs. The top line shows characteristics of mAbs with (red) and without (blue) RG motifs in CDRH3s. The bottom line shows characteristics of mAbs with (red) and without (blue) WD motifs in CDRL3s. The left column shows the numbers of mAbs with and without CDR3 motifs with respect to two most common V genes: IGHV1-46 and IGHV3-23 for RG motifs in CDRH3, IGKV3-20 and IGLV1-51 for WD motifs in CDRL3s. The middle and right columns show the responses of iGL and mature mAbs to stem peptides of SARS-CoV-2 and MERS-CoV, respectively. P-values of associations between RG / WD motifs and the responses are shown on tops of the plots and denoted as follows: ns≥0.05, *<0.05, **<0.005. P-values are computed using linear regression. **c.** ELISA-based epitope mapping of S2 stem-helix bnAbs with SARS-CoV-2 stem alanine scan peptides (25mer). Heatmap shows fold-changes in EC_50_ binding titers of mAb binding to SARS-CoV-2 stem-helix peptide alanine mutants compared with the WT peptide. SARS-CoV-2 stem-helix residue positions targeted (2-fold or higher decrease in EC_50_ binding titer compared to WT stem peptide) is indicated in colors. Three hydrophobic residues, F^1148^, L^1152^ and F^1156^, were commonly targeted by stem-helix bnAbs and that form the core of the bnAb epitope. Association of dependence on the stem bnAb core epitope residues with heavy (IGHV1-46 and IGHV3-23) and light (IGKV3-20 and IGLV1-51) chain genes usage and CDRL3 lengths is shown. **d-g.** A SARS-CoV-2 spike protein cartoon depicts the S2-stem epitope region in green at the base of the prefusion spike ectodomain **(d)**. Sequence conservation of stem-helix hydrophobic core epitope residues (F^1148^, L^1152^ and F^1156^) across β-coronavirus spikes (PDB: 6XR8) **(e)**. D^1146^ stem-helix residue is also indicated. Side **(f)** and top **(g)** views of spike stem-helix region highlight the core epitope residues. **h.i.** Interactions between SARS-CoV-2 S2 stem helix with **(h)** S2P6 and **(i)** CC40.8 highlighting the contribution of antibody germline-encoded residues in recognition of hydrophobic stem-helix core epitope. The SARS-CoV-2 S2 stem helices are shown in green, while heavy and light chains of antibodies in orange and yellow, respectively. Germline gene encoded residues are highlighted in red. Structures with PDB codes 7RNJ and 7SJS are used for S2P6 and CC40.8, respectively.

In contrast to binding, neutralization of SARS-CoV-2 and MERS-CoV by stem-helix bnAb iGLs was absent (Supplementary Fig. 5d). The result suggests that, although overall SHM levels do not correlate with binding or neutralization, key antibody mutations are critical for the neutralization phenotype to attain sufficient affinity for neutralization to be observed.

Altogether, we have isolated a large panel of human β-CoV bnAbs that are enriched in certain germline gene features suggesting the potential value of a highly targeted approach (*53, 54, 63*) to induce pan-betacoronavirus bnAbs by vaccines in which the immunogen and vaccination strategies are appropriately designed.

### Spike stem-helix bnAbs recognize a common hydrophobic core epitope

To determine the epitope specificities of the isolated stem-helix bnAbs and potential association with antibody immunogenetic properties, we performed binding of all 32 stem bnAbs to alanine scanning mutants of the SARS-CoV-2 stem-peptide (Fig. 3c, Supplementary Fig. 7). A dependence on three hydrophobic residues, F^1148^, L^1152^, and F^1156^, by many bnAbs that form a common core epitope was identified but the relative dependence of bnAb lineages on each of the hydrophobic core residues varied. Many of the IGHV1-46-encoded bnAbs were paired with IGVK3-20 or IGLV1-51 light chain and all except two bnAbs possessed a CDRL3 of 11 residues. The IGHV3-23-encoded bnAbs showed dependence on 1 or 2 hydrophobic core epitope residue and some lineages showed dependence on an upstream acidic residue, D^1146^. All of the IGHV3-23 encoded bnAbs were paired with a IGVK3-20 light chain with a 9-residue long CDRL3 loop. The non −IGHV1-46 or −IGHV3-23-encoded stem-helix bnAbs were also dependent on one or more hydrophobic core epitope residues with one exception. Structural analysis of the IGHV1-46-encoded S2P6 or IGHV3-23-encoded CC40.8 stem-helix bnAbs shows that antibody germline gene-encoded residues are involved in recognition of the hydrophobic bnAb epitope (Fig. 3d-i). Overall, hydrophobic core residues in the spike fusion machinery, which are highly conserved across betacoronaviruses, are important targets for S2 bnAbs. Notably, the hydrophobic core epitope residues on the pre-fusion S-trimer are poorly accessible and partial disruption of the stem-helix region may be needed to favorably expose this bnAb site to engage desired B cell responses (*43-45, 48*).

### Stem-helix bnAbs protect against challenge with diverse β-CoVs

To determine the protective efficacy of the stem-helix bnAbs, we prophylactically treated aged mice (*65*) with individual antibodies followed by virus challenge. We selected two of the broadest and potent stem-helix bnAbs, CC68.109 and CC99.103, and investigated their *in vivo* protective efficacy against all three major human disease-causing betacoronaviruses; SARS-CoV-2, SARS-CoV-1 and MERS-CoV. Prior to the challenge experiments, we examined neutralization of SARS-CoV-2 and MERS-CoV replication-competent viruses by the two candidate bnAbs and compared with that of pseudoviruses (Supplementary Fig. 8). The neutralization IC_50_s of the stem-helix bnAbs were comparable for SARS-CoV-2 across the two assay formats while the titers with replication-competent MERS-CoV were more effective (lower IC_50_ values) compared to the pseudovirus format. The two stem-helix bnAbs, individually, or a DEN3 control antibody were administered intra-peritoneally (i.p.) at 300μg/animal into 9 groups of 10 animals (3 groups per antibody; Fig. 4a). 12h prior to the virus challenge, the test antibody in each animal group was administered followed by intranasal (i.n.) challenge with one of three mouse-adapted (MA) betacoronaviruses, (MA10-SARS-2 = SARS-CoV-2; MA15-SARS-1 = SARS-CoV-1 or M35c4-MERS = MERS-CoV) (Fig. 4a) (*65-67*). Post virus challenge, the animals were monitored for signs of clinical disease due to infection, including daily weight changes, and pulmonary function. Animals were euthanized at day 2 or day 4 post infection and lung tissues were harvested to assess gross pathology. Compared to the control antibody DEN3-treated animal groups, the stem-helix bnAb-treated animals in all three betacoronaviruses challenge experiments showed substantially reduced weight loss (Fig. 4b, e, h), reduced hemorrhage (Fig. 4c, f, i), and normal pulmonary function (Fig. 4d, g, j), suggesting a protective role for the bnAbs.

**Figure. 4.**
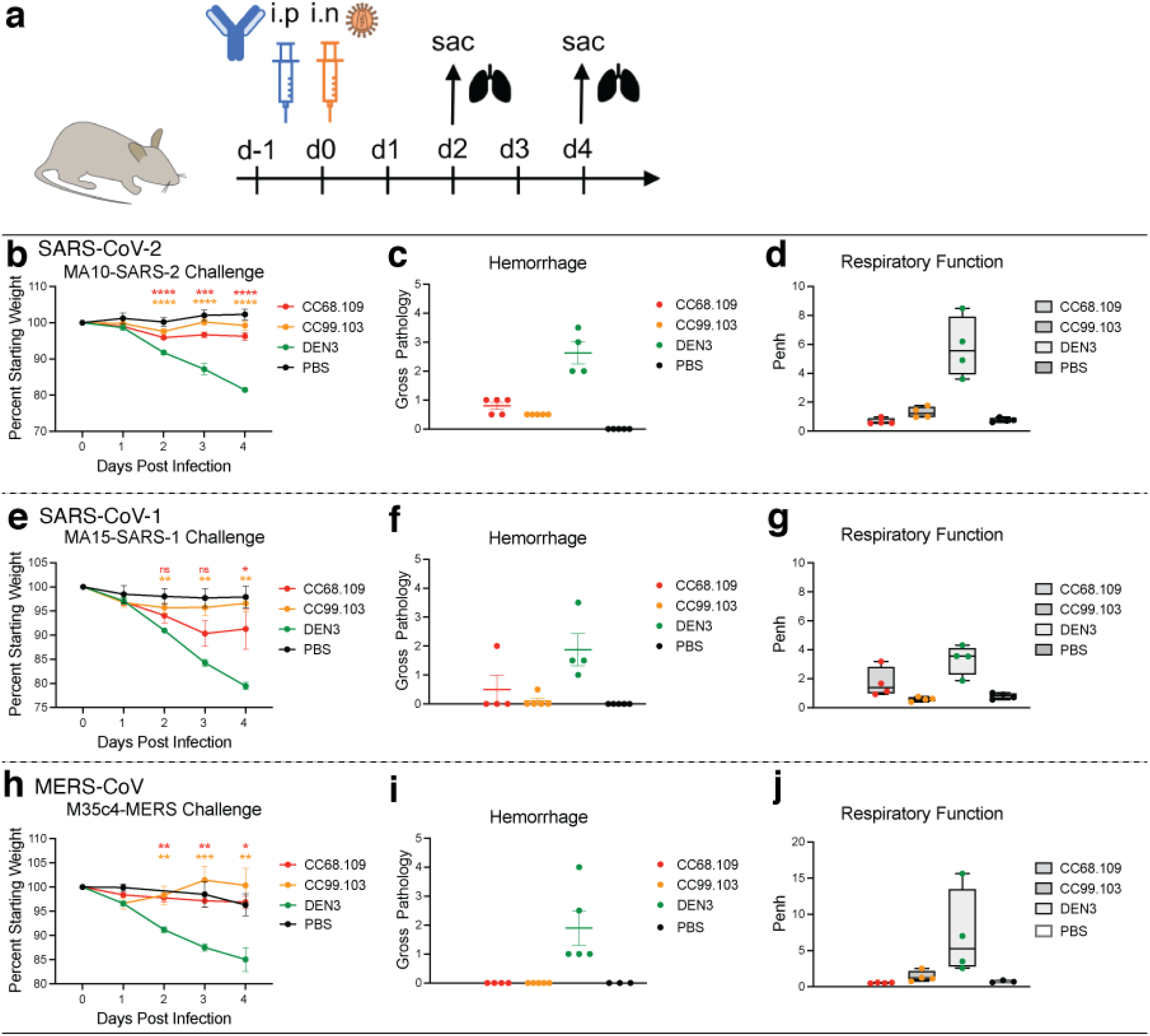
Prophylactic treatment of aged mice with S2 stem-helix bnAbs protects against challenge with diverse betacoronaviruses. **a.** Two S2 stem-helix bnAbs (CC68.109 and CC99.103) individually, or a DEN3 control antibody were administered intra-peritoneally (i.p.) at 300μg per animal into 9 groups of aged mice (10 animals per group). Each group of animals was challenged intra-nasally (i.n.) 12h after antibody infusion with one of 3 mouse-adapted (MA) betacoronaviruses, (MA10-SARS-2 = SARS-CoV-2; 1 × 10^3^ plaque forming units (PFU), MA15-SARS1 = SARS-CoV-1; 1 × 10^5^ PFU or M35c4-MERS = MERS-CoV; 1 × 10^3^ PFU). As a control, groups of mice were exposed only to PBS in the absence of virus**. b., e., h.** Percent weight change in S2 stem-helix bnAbs or DEN3 control antibody-treated animals after challenge with mouse-adapted betacoronaviruses. Percent weight change was calculated from day 0 starting weight for all animals. **c., f., i.** Day 2 post-infection Hemorrhage (Gross Pathology score) scored at tissue harvest in mice prophylactically treated with S2 stem-helix bnAbs or DEN3 control mAb. **d., g., j.** Day 2 post-infection pulmonary function (shown as Penh score) was measured by whole body plethysmography in mice prophylactically treated with S2 stem-helix bnAbs or DEN3 control mAb. Statistical comparisons between groups were performed using a Kruskal-Wallis non-parametric test and significance was calculated with Dunnett's multiple comparisons test between each experimental group and the DEN3 control Ab group. (p <0.05, **p <0.01, ***p <0.001; ****p < 0.0001; ns- p >0.05).

Overall, both stem-helix bnAbs protected against severe betacoronavirus disease, CC99.103 being slightly more protective than CC68.109 bnAb.

## Discussion

In terms of passive antibody treatment, the ability of single mAbs to protect against the two SARS viruses and MERS in the small animal model is encouraging for their potential adoption as stockpiled reagents to tackle future outbreaks of viral infection, including novel related betacoronaviruses. Prophylaxis or treatment very early in infection is more likely to be successful than therapy once symptoms are established, based on experience with SARS-CoV-2 (*68, 69*). The use of S2 bnAbs, possibly in a cocktail with the most appropriate RBD bnAbs, may be the optimal approach for SARS-CoV-2 prophylaxis, especially as new variants such as Omicron emerge. The dose of S2 bnAbs required to be effective, given the typically lower neutralization potencies of such nAbs compared to RBD nAbs, may be an issue for translation. However, the studies in animal models (*44, 45*) suggest that S2 bnAbs protect at much lower serum concentrations than would be predicted by their IC_50_ neutralizing titers-i.e. they “punch above their weight”. Therefore, in addition to neutralization, effector functions of the S2 bnAbs may be important for protection (*45, 70-72*) and clinical studies will be required to investigate this phenomenon in humans.

In terms of vaccine design, a rational strategy is strongly favored by the availability of a panel of bnAbs rather than single mAbs so that the broadly neutralizing epitope can be more precisely defined and the qualities of nAbs required for broad neutralization determined (*53, 54, 56, 63, 73, 74*). Here, we identified critical hydrophobic S2 residues involved in bnAb binding and showed the prevalence of a IGVH1-46/IGVK3-20 antibody pairing with restricted CDRH3 and CDRL3 lengths in S2 bnAbs. Accordingly, rational vaccine design strategies may take advantage of these germline gene features to develop immunogens that can induce protective antibody responses to this site (*52-54, 56, 63*). Accessibility of the S2 stem-helix bnAb site on spike immunogen to effectively engage desired B cell responses might be challenging. Nevertheless, approaches to scaffold immunogen designs that can accommodate these features are now available to be deployed (*46, 52, 56, 75-77*).

In summary, we isolated the largest panel of β-CoV bnAbs to date and revealed the molecular basis for their broad protection. The bnAbs provide a detailed framework for rational design of broad coronavirus vaccines and themselves could be used as reagents to counter betacoronavirus spillovers.

## Author contributions

P.Z., G.S., R.S.B., L.E.G., D.R.B. and R.A. conceived and designed the study. N.B., M.P., E.G., S.A.R., D.M.S., and T.F.R. recruited donors and collected and processed plasma and PBMC samples. P.Z., G.S., W.H., R.M., K.D., S.C., P.Y., L.P., T.C., O.L. and F.A. performed BLI, ELISA, virus preparation, neutralization and isolation and characterization of monoclonal antibodies. Y.S. performed immunogenetic analysis of antibodies. M.Y. and H.L. generated antibody-antigen structural models. L.V.T. performed live virus neutralizations assays and D.R.M., A.S., and L.E.G. conducted *in vivo* animal protection studies. P.Z., G.S., W.H., N.B., L.V.T., D.R.M., A.S., F.A., P.Y., L.P., K.D., R.M., S.C., T.C., M.Y., H.L., O.L., M.P., E.G., D.N., J.G.J., I.A.W., Y.S., T.F.R., R.S.B., L.E.G., D.R.B. and R.A. designed the experiments and/or analyzed the data. P.Z., G.S., D.R.B. and R.A. wrote the paper, and all authors reviewed and edited the paper.

## Acknowledgements

We thank all the human cohort participants for donating samples. This work was supported by NIH CHAVD UM1 AI44462 (D.R.B.), NIH R61 AI161818 (R.A.), the IAVI Neutralizing Antibody Center, the Bill and Melinda Gates Foundation INV-004923 (I.A.W., D.R.B.), the Translational Virology Core of the San Diego Center for AIDS Research (CFAR) grant NIH AI036214 (D.M.S.), NIH 5T32AI007384 (S.A.R.), NIH U54 CA260543 and AI157155 (R.S.B), NIH R21 AI145372 (L.E.G.), and the John and Mary Tu Foundation and the James B. Pendleton Charitable Trust (D.M.S. and D.R.B.). L.V.T. is supported by Pfizer NCBiotech Distinguished Postdoctoral Fellowship in Gene Therapy. D.R.M. is currently supported by a Burroughs Wellcome Fund Postdoctoral Enrichment Program Award and a Hanna H. Gray Fellowship from the Howard Hugues Medical Institute.

## Data availability

The data supporting the findings of this study are available within the paper and its supplementary information files or from the corresponding author upon reasonable request. Antibody sequences have been deposited in GenBank under accession numbers XXXX-XXXX. Antibody plasmids are available from Raiees Andrabi or Dennis Burton under an MTA from The Scripps Research Institute.

## Methods

### COVID-19 infected-vaccinated donors

Sera and PBMC samples from convalescent COVID-19 donors, vaccinated donors, and COVID-19-recovered vaccinated donors, were provided through the “Collection of Biospecimens from Persons Under Investigation for 2019-Novel Coronavirus Infection to Understand Viral Shedding and Immune Response Study” UCSD IRB# 200236 as reported earlier (*35*). The protocol was approved by the UCSD Human Research Protection Program. Convalescent donor samples were collected based on COVID-19 diagnosis regardless of gender, race, ethnicity, disease severity, or other medical conditions. All human donors were assessed for medical decision-making capacity using a standardized, approved assessment, and voluntarily gave informed consent prior to being enrolled in the study.

### Plasmid construction

To generate soluble S ectodomain proteins from SARS-CoV-1 (residues 1-1190; GenBank: AAP13567) ,SARS-CoV-2 (residues 1-1208; GenBank: MN908947), HCoV-HKU1 (residue 1-1295; GenBank: YP_173238.1), HCoV-OC43 (residues 1-1300; GenBank: AAX84792.1), MERS-CoV (residues 1-1291; GenBank: APB87319.1), HCoV-229E (residues 1-1110; GenBank: NP_073551.1) and HCoV-NL63 (residues 1-1291; GenBank: YP_003767.1), we synthesized the DNA fragments from GeneArt (Life Technologies) and cloned them into the phCMV3 vector (Genlantis cat.# P003300). In order to produce the stable trimeric prefusion spike proteins, double proline substitutions (2P) were introduced into the S2 subunit: K968/V969 in SARS-CoV-1, K986/V987 in SARS-CoV-2, V1060/L1061 in MERS-CoV, A1071/L1072 in HCoV-HKU1, A1078/L1079 in HCoV-OC43, S1052/I1053 in HCoV-NL63 and T871/I872 in HCoV-229E were replaced by proline. The furin cleavage sites (in SARS-CoV-2 residues 682–685, in SARS-CoV-1 residues 664–667, in HCoV-HKU1 residues 756-760, in HCoV-OC43 residues 762–766, in MERS-CoV residues 748–751, in HCoV-229E residues 564–567 and in HCoV-NL63 residues 745–748) were replaced by a “GSAS” linker; the trimerization T4 fibritin motif was incorporated at the C-terminus of the S proteins. To purify and biotinylate the spike proteins, the HRV-3C protease cleavage site, 6x HisTag, and AviTag spaced by GS-linkers were added to the C-terminus after the trimerization motif. To generate pseudoviruses of MERS-CoV and sarbecoviruses, the DNA fragments encoding the spikes of MERS-CoV and sarbecoviruses without the ER retrieval signal were codon-optimized and synthesized at GeneArt (Life Technologies). The spike encoding genes of Pang17 (residues 1-1249, GenBank: QIA48632.1), WIV1 (residues 1-1238, GenBank: KF367457) and SHC014 (residue 1-1238, GenBank: AGZ48806.1) were constructed into the phCMV3 vector (Genlantis cat.# P003300) using the Gibson assembly (New England Biolabs, cat.# E2621L) according to the manufacturer’s instructions.

### Cell lines

FreeStyle293-F cells (Thermo Fisher Scientific cat.# R79007) were grown in FreeStyl 293 Expression Medium (Gibco cat.# 12338018), and Expi293F cells (Gibco cat.# A14527) were maintained in Expi293 Expression Medium (Gibco cat.# A1435101). Suspension cells were incubated in the shaker at 150 rpm, 37°C, 8% CO_2_. Adherent HEK293T cells and HeLa-ACE2 cells were grown in Dulbecco's Modified Eagle Medium (DMEM) with 10% heat-inactivated FBS, 4mM L-Glutamine and 1% P/S, maintaining in the incubator at 37°C, 5% CO_2_. The stable hACE2 or hDPP4-expressing HeLa cell line was generated using an ACE2 lentivirus protocol previously described (*7*). Briefly, the pBOB-hACE2 or hDPP4 plasmid and lentiviral packaging plasmids (pMDL, pREV, and pVSV-G (Addgene #12251, #12253, #8454)) were co-transfected into HeLa cells using Lipofectamine 2000 reagent (ThermoFisher Scientific cat.# 11668019).

### Expression and purification of HCoV S-proteins

To express the soluble human coronavirus (HCoV) S ectodomain proteins with His-tag or with both His- and Avi-tag at the C-terminus, 350 μg plasmids in 15ml Opti-MEM™ (Thermo Fisher Scientific cat.# 31985070) were filtered and mixed with 1.8 ml 40K PEI (1mg/ml) in 15ml Opti-MEM™, then incubated at room temperature for 30 min and transferred into 1L FreeStyle293-F cells at the density of 1 million cells/ml. Four days after transfection, the cell cultures were centrifuged at 2500xg for 15 min and filtered through 0.22μm membrane. The His-tagged proteins were purified with the HisPur Ni-NTA Resin (Thermo Fisher Scientific cat.# 88221). After washing by wash buffer (25 mM Imidazole, pH 7.4) for at least 3 bed volumes, the protein was eluted with 25 ml elution buffer (250 mM Imidazole, pH 7.4) at slow gravity speed (~4 sec/drop), then was buffer exchanged into PBS and concentrated using 100K Amicon tubes (Millipore cat.# UFC910024). After being further purified by size-exclusion chromatography by Superdex 200 Increase 10/300 GL column (GE Healthcare cat.# GE28-9909-44), the protein was pooled and concentrated again for further use.

### Flow cytometry B cell profiling and monoclonal antibody isolation

Flow cytometry of PBMC samples from infected-vaccinated human donors were conducted following methods described previously (*7*). 10ml RPMI1640 medium (Thermo Fisher Scientific, cat.# 11875085) with 50% FBS was pre-warmed to 37°C and used to thaw the frozen PBMC samples, followed by centrifugation at 400xg for 5 min. After discarding supernatant, the cells were resuspended in a 5 ml FACS buffer (PBS, 2% FBS, 2 mM EDTA). Fluorescently labeled antibodies specific for cell surface markers were prepared as 1:100 dilution as a master mix in FACS buffer, to stain the PBMC samples for CD3 (APC-Cy7, BD Pharmingen cat.# 557757), CD4 (APC-Cy7, Biolegend cat.# 317418), CD8 (APC-Cy7, BD Pharmingen cat.# 557760), CD14 (APC-H7, BD Pharmingen cat.# 561384), CD19 (PerCP-Cy5.5, Fisher Scientific cat.# NC9963455), CD20 (PerCP-Cy5.5, Biolegend, cat.# 302326), IgG (BV605, BD Pharmingen cat.# 563246) and IgM (PE, Biolegend, cat.# 314508). Meanwhile, SARS-CoV-2 S protein with Avi-tag was conjugated to streptavidin-BV421 (BD Pharmingen cat.# 563259) and streptavidin-AF488 (Invitrogen cat.# S11223), respectively, and the MERS-CoV S protein with Avi-tag was conjugated to streptavidin-AF647 (Invitrogen cat.# S21374). After incubating the cells with Ab mixture for cell surface markers for 15 min in dark, S protein-probes were added to the samples and incubated on ice in the dark for 30 min. FVS510 Live/Dead stain (Thermo Fisher Scientific cat.# L34966) in FACS buffer (1:300) was then added to the samples and incubated on ice in the dark for 15 min. After washing with FACS buffer, the stained cells were resuspended in 500 μl of FACS buffer per 10-20 million cells, filtered through the 70-μm mesh cap into FACS tubes (Fisher Scientific cat.# 08-771-23) and sorted for S protein-specific memory B cells using BD FACSMelody sorter. In brief, after gating of lymphocytes (SSC-A vs. FSC-A) and singlets (SSC-W vs SSC-H and FSC-H vs. FSC-W), live cells were identified by the negative FVS510 live/dead staining phenotype. The CD3^−^ CD4^−^ CD8^−^ CD14^−^ CD19^+^ CD20^+^ cells were gated as B cells. By selecting the IgG^+^ IgM-population, the cells were sequentially gated for SARS-CoV-2-S-BV421^+^ SARS-CoV-2-S-AF488^+^ MERS-CoV-S-AF647^+^ reactivity. Triple positive memory B cells was sorted as single cells into 96-well plates on a cooling platform. Superscript IV Reverse Transcriptase (Invitrogen cat.# 18090010), 10mM dNTPs (Invitrogen cat.# 18427088), random hexamers (Gene Link cat.# 26-4000-03), Ig gene-specific primers, 0.1M DTT, RNAseOUT (Invitrogen cat.# 10777019), and 10% Igepal (Sigma-Aldrich cat.# 18896) were used in the reverse transcription PCR reaction to generate cDNA from the sorted cells right after sorting. Hot Start DNA Polymerases ((QIAGEN cat.# 203643) and specific primer sets described previously (*78, 79*) were used to perform two rounds of nested PCR reactions to amplify IgG heavy and light chain variable regions using cDNAs as template. After being purified with SPRI beads according to manufacturer’s instructions (Beckman Coulter cat.# B23318), PCR products were constructed into expression vectors encoding human IgG1 or Ig kappa/lambda constant domains, respectively, by Gibson assembly (New England Biolabs cat.# E2621L), then transformed into competent *E.coli* cells. Single colonies were picked for sequencing and analysis on IMGT V-Quest online tool (http://www.imgt.org) and downstream plasmid production.

### Expression and purification of monoclonal antibodies

Plasmids of the paired heavy and light chains generated after sorting were co-transfected into Expi293F cells to produce monoclonal antibodies. Briefly, 12μg heavy chain plasmid and 12 μg of light chain plasmid were added into 3ml of Opti-MEM™ (Thermo Fisher Scientific cat.# 31985070), after inverting, 24μl of FectoPRO (Polyplus cat.# #116-001) reagent was added into the mixture and inverted. Incubation at room temperature for 10min was done before adding the mixture into 30ml of Expi293F cells at 2.8 million cells/ml and incubating in the shaker. 24 hours after transfection, 300μl of 300mM sodium valproic acid solution and 275μl of 45% Glucose solution was used to feed each cell culture. Four days post transfection, supernatants of cell cultures were collected by centrifugation at 2500xg for 15 min and filtering through 0.22μm membrane. Protein A Sepharose (GE Healthcare cat.# 45002982) and Protein G Sepharose (GE Healthcare cat.# 45000118) were mixed at 1:1 ratio before adding into the supernatant and rotating overnight at 4°C. The solution was then loaded into Econo-Pac columns (BioRad cat.# 7321010), washed with 1 column volume of PBS, and antibodies were eluted with 10ml of 0.2 M citric acid (pH 2.67). The elution was collected into a tube containing 1ml of 2M Tris Base solution. 30K Amicon centrifugal filters (Millipore cat.# UFC903024) were used for buffer exchange into PBS and further concentrating into smaller volumes.

### ELISA using peptides or recombinant proteins

N-terminal biotinylated peptides corresponding to stem helix of SARS-CoV-1/2, MERS-CoV, HCoV-HKU1, HCoV-OC43, HCoV-229E and HCoV-NL63 were synthesized at A&A Labs (Synthetic Biomolecules) (*44*). For peptide ELISA, streptavidin (Jackson Immuno Research Labs cat.# 016-000-084) was coated at 2 μg/ml in PBS onto 96-well half-area high binding plates (Corning, 3690) overnight at 4°C. For recombinant protein ELISA, mouse anti-His antibody (Invitrogen cat. # MA1-21315-1MG) was used at the same concentration to coat the plates. After washing by 0.05% PBST 3 times, 3% BSA was used to block the plates for 2h at 37°C. Then 1 μg/ml of N-terminal biotinylated peptide or 2 μg/ml of His-tagged recombinant spike proteins were applied to plates and incubated for 1h at RT. After washing by 0.05% PBST 3 times, serially diluted serum samples or antibodies were added into plates and incubated for 1h at RT. After another washing, alkaline phosphatase-conjugated goat anti-human IgG Fc secondary antibody (Jackson ImmunoResearch cat.# 109-055-008) was added in 1:1000 dilution and incubated for 1h at RT. After the final wash, phosphatase substrate (Sigma-Aldrich cat.# S0942-200TAB) dissolved in staining buffer was added into each well. Absorption was measured at 405 nm. Fifty percent maximal response concentrations (EC50) were calculated using the Asymmetrical dose-response model of Richard’s version in GraphPad Prism 7 (GraphPad Software). To identify critical residues for antibody binding, single alanine mutations were introduced onto the 25-mer stem helix peptide that comprises the linear epitope. These peptides were synthesized at A&A Labs (Synthetic Biomolecules). ELISA as described above was used to test antibody reactivity against peptides with single alanine substitutions.

### Pseudovirus production

HIV-based lentivirus backbone plasmid pCMV-dR8.2 dvpr (Addgene #8455), pBOB-Luciferase (Addgene #170674) were co-transfected into HEK293T cells along with full-length or variously truncated SARS-CoV1, WIV1, SHC014, Pang17, SARS-COV2, SARS-CoV-2 variants of concern [(B.1.1.7(alpha), B.1.351 (beta), P.1 (gamma), B.1.617.2 (delta) and B.1.1.529 (Omicron)] and MERS-CoV spike using Lipofectamine 2000 (ThermoFisher Scientific cat.# 11668019) to produce single-round infection-competent pseudoviruses (*80*). The medium was changed 12-16 hours post transfection. Pseudovirus-containing supernatants were collected 48 hours post transfection and the viral titers were measured by luciferase activity in relative light units (RLU) (Bright-Glo Luciferase Assay System, Promega cat.# E2620). The supernatants were aliquoted and stored at −80°C until further use.

### Neutralization assay

Pseudotyped viral neutralization assay was performed as previously reported (*7*). In brief, neutralization assays were performed by adding 25μl of pseudovirus into 25μl serial dilutions of purified antibodies or plasma from human donors, the mixture was then dispensed into a 96-well plate incubated for one hour at 37°C, then 10,000 HeLa-hACE2 or hDPP4 cells/ well (in 50μl of media containing 20μg/ml Dextran) were directly added to the mixture. After incubation at 37°C for 42-48 h, luciferase activity was measured. Neutralizing activity was measured by reduction in luciferase activity compared to the virus controls. Fifty percent maximal inhibitory concentrations (IC_50_), the concentrations required to inhibit infection by 50% compared to the controls, were calculated using the dose-response-inhibition model with 5-parameter Hill slope equation in GraphPad Prism 7 (GraphPad Software).

### Neutralization Assay of Replication Competent CoVs

Vero E6 cells (ATCC-C1008) were seeded at 2×10^4^ cells/well in a black-well, black-wall, tissue culture treated, 96-well plate (Corning Cat. #3916) 24 h before the assay. Abs were diluted in MEM supplemented with 5%FBS and 1%Pen/Strep media to obtain an 8-point, 3-fold dilution curve with starting concentration at 20 μg/ml. Eight hundred Pfu of SARS2-nLuc and MERS-nLuc replication competent viruses were mixed with Abs at a 1:1 ratio and incubated at 37°C for 1 h. One-hundred microliters of virus and Ab mix was added to each well and incubated at 37ºC + 5% CO_2_ for 20 to 22 h. Luciferase activities were measured by the Nano-Glo Luciferase Assay System (Promega Cat. #N1130) following the manufacturer’s protocol using a GloMax luminometer (Promega). Percent inhibition and IC_50_ were calculated as pseudovirus neutralization assay described above. All experiments were performed as duplicate and independent repeated for three times. All the live virus experiments were performed under biosafety level 3 (BSL-3) conditions at negative pressure, by operators in Tyvek suits wearing personal powered-air purifying respirators.

### HEp2 epithelial cell polyreactive assay

According to manufacturer’s instructions, HEp2 slides (Hemagen cat.# 902360) were used to determine the reactivity of monoclonal antibodies to human epithelial type 2 (HEp2) by indirect immunofluorescence. Briefly, monoclonal antibody was diluted into 50μg/ml by PBS and then added onto immobilized HEp2 slides and incubated for 30 min at RT. After washing by PBS for 3 times, one drop of FITC-conjugated goat anti-human IgG was added onto each well and incubated in the dark for 30 min at RT. After washing, the coverslip was added to HEp2 slide with glycerol and the images were photographed on a Nikon fluorescence microscope for FITC detection.

### Polyspecificity reagent (PSR) ELISA

Solubilized CHO cell membrane protein (SMP), human insulin (Sigma-Aldrich cat.# I2643), single strand DNA (Sigma-Aldrich cat.# D8899) were coated onto 96-well half-area high-binding plates (Corning cat.# 3690) at 5μg /ml in PBS overnight at 4°C. After washing with PBST, plates were blocked with 3% BSA for 2h at 37°C. Antibody samples were diluted at 50μg /ml in 1% BSA with 5-fold serial dilution and then added in plates to incubate for 1h at room temperature (*7*). The assay was performed as described in section “ELISA using peptides or recombinant proteins”.

### CELISA binding

Flow cytometry-based Cell-ELISA (CELISA) binding of mAbs with HCoV spikes was performed as described previously (*43, 81*). A total of 4×10^6^ HEK293T cells were seeded into 10cm round cell culture dishes and incubated at 37°C. After 24h, HEK293T cells were transfected with plasmids encoding full-length HCoV spikes and were incubated for 36-48h at 37°C. The cells were harvested and distributed into 96-well round-bottom tissue culture plates for individual staining reactions. For each staining reaction, cells were washed three times with 200μl FACS buffer (1xPBS, 2%FBS, 1mM EDTA). The cells were stained for 1h on ice in 50μl staining buffer with 10μg/ml of primary antibody. After washing three times with 200μl FACS buffer, the cells were stained with 50μl/well of 1:200 diluted R-phycoerythrin (PE)-conjugated mouse anti-human IgG Fc antibody (SouthernBiotech cat.# 9040-09) and 1:1000 dilution of Zombie-NIR viability dye (BioLegend cat.# 423105) on ice in dark for 45min. Following three washes with FACS buffer, the cells were resuspended and analyzed by flow cytometry (BD Lyrics cytometer), and the binding data were generated by calculating the Mean Fluorescence Intensity using FlowJo 10 software. Mock-transfected 293T cells were used as a negative control.

### BioLayer Interferometry binding (BLI)

Octet K2 system (ForteBio) was used to determine the monoclonal antibody binding with S-proteins or selected peptides. IgG was first captured for 60s by anti-human IgG Fc capture (AHC) biosensors (ForteBio cat.# 18-5063), then baseline was provided in Octet buffer (PBS with 0.1% Tween) for another 60s. After that, the sensors were transferred into wells containing diluted HCoV S-proteins for 120s for association, and into Octet buffer for disassociation for 240s. Selected peptides that were N-terminal biotinylated were diluted in Octet buffer and first captured for 60s by the hydrated streptavidin biosensors (ForteBio cat.# 18-5020), then unbound peptides were removed by transferring into Octet buffer for 60s to provide the baseline. Then the sensors were immersed into monoclonal antibodies in Octet buffer for 120s for association, followed by transferring into Octet buffer for 240s for dissociation. The data generated were analyzed using the ForteBio Data Analysis software for correction, and the kinetic curves were fit to 1:1 binding mode. Note that the IgG: spike protomer binding can be a mixed population of 2:1 and 1:1, such that the term ‘apparent affinity’ dissociation constants (K_D_^App^) are shown to reflect the binding affinity between IgGs and spike trimers tested.

### Antibody immunogenetics analysis

Heavy and light chain sequences of mature antibodies were processed using DiversityAnalyzer tool (*82*). For each CDRH3 translated in the amino acid alphabet, all its *k*-mers were extracted, where *k*=2, 3, 4. *K*-mers appearing in at least 20% of HCDR3s were reported as motifs. In total 10 motifs were reported for CDRH3s: AR, ARG, AS, DY, FD, FDY, GS, GV, RG, SS. Each heavy chain sequence was labeled by whether its CDRH3 contains a given motif. The same procedure was applied to CDRL3s and reported 16 motifs: DS, DSS, FT, GS, PP, QQ, QQY, QY, QYG, SP, SPP, SS, SSP, SSPP, WD, YG. For each CDRH3 motif, the linear regression model was applied to estimate the impact of the motif presence (denoted as “yes” or “no”) and the type of antibody (denoted as “iGL” or “mature”) on the responses of 32 mature antibodies to the stem helix peptides of SARS-CoV-2 and MERS-CoV viruses. The same method was applied to estimate the impact of the presence of LCDR3 motifs. Heavy and light chain sequences of the same antibody were concatenated into a single sequence and collected across all 32 antibodies. The phylogenetic tree derived from the concatenated sequences was constructed using ClusterW2 tool (*83*) and visualized using the Iroki tool (*84*).

### *In vivo* virus challenge in mouse model

All mouse experiments were performed at the University of North Carolina, NIH/PHS Animal Welfare Assurance Number: D16-00256 (A3410-01), under approved IACUC protocols. The animal manipulation and virus work was performed in a Class 2A biological safety cabinet in a BSL3 approved facility and workers wore PAPRs, tyvek suites and were double gloved. 12-month-old female Balb/c mice (strain 047) were purchased from Envigo for Sarbecovirus challenge experiments (*65, 85*). C57Bl/6 288/330+/+ mice, which encode two human codons in the mouse dipeptidyl peptidase gene, were used for MERS-CoV mouse adapted challenge experiments (*66*). Mice were housed in individually ventilated Seal-Safe cages, provided food and water *ad libitum* and allowed to acclimate at least seven days before experimental use. Twelve hours prior to infection, 300μg antibody was injected into mice intraperitoneally. Immediately prior to infection, mice were anesthetized by injection of ketamine and xylazine intraperitoneally and weighed. Virus (SARS-CoV MA15, SARS-CoV2 MA10 and mouse adapted MERS-CoV-M35c4) was diluted in 50μl sterile PBS and administered intranasally (*65-67, 85*). Mice were weighed daily and observed for signs of disease. The mice were euthanized via isoflurane overdose at the designated timepoint, followed by assessment of gross lung pathology and collection of the inferior lobe for virus titration. Respiratory function was measured at day2 post infection via Buxco whole body plethysmography, as previously described (*86*).

### Virus titration

SARS-CoV-2-MA10, SARS-CoV-1-MA15 and MERS-CoV-M35c4 were grown and titered using VeroE6 cells as previously described (*87*). Briefly, lung tissue was homogenized in 1ml sterile PBS via Magnalyser (Roche), centrifuged to pellet debris, plated in 10-fold serial dilutions on VeroE6 cells on a 6-well plate and covered with a 1:1 mixture of 1.6% agarose and media. At two (SARS-CoV-1) or three (SARS-CoV-2) days post plating, cells were stained with neutral red and plaques counted.

### Statistical Analysis

Statistical analysis was performed using Graph Pad Prism 8, Graph Pad Software, San Diego, California, USA. ID_50_ or IC_50_ titers were compared using the non-parametric unpaired Mann-Whitney-U test. The correlation between two groups was determined by Spearman rank test. Groups of data were compared using the Kruskal-Wallis non-parametric test. Dunnett's multiple comparisons test were also performed between experimental groups. Data were considered statistically significant at p < 0.05.

---

**Supplementary Figure 1.**
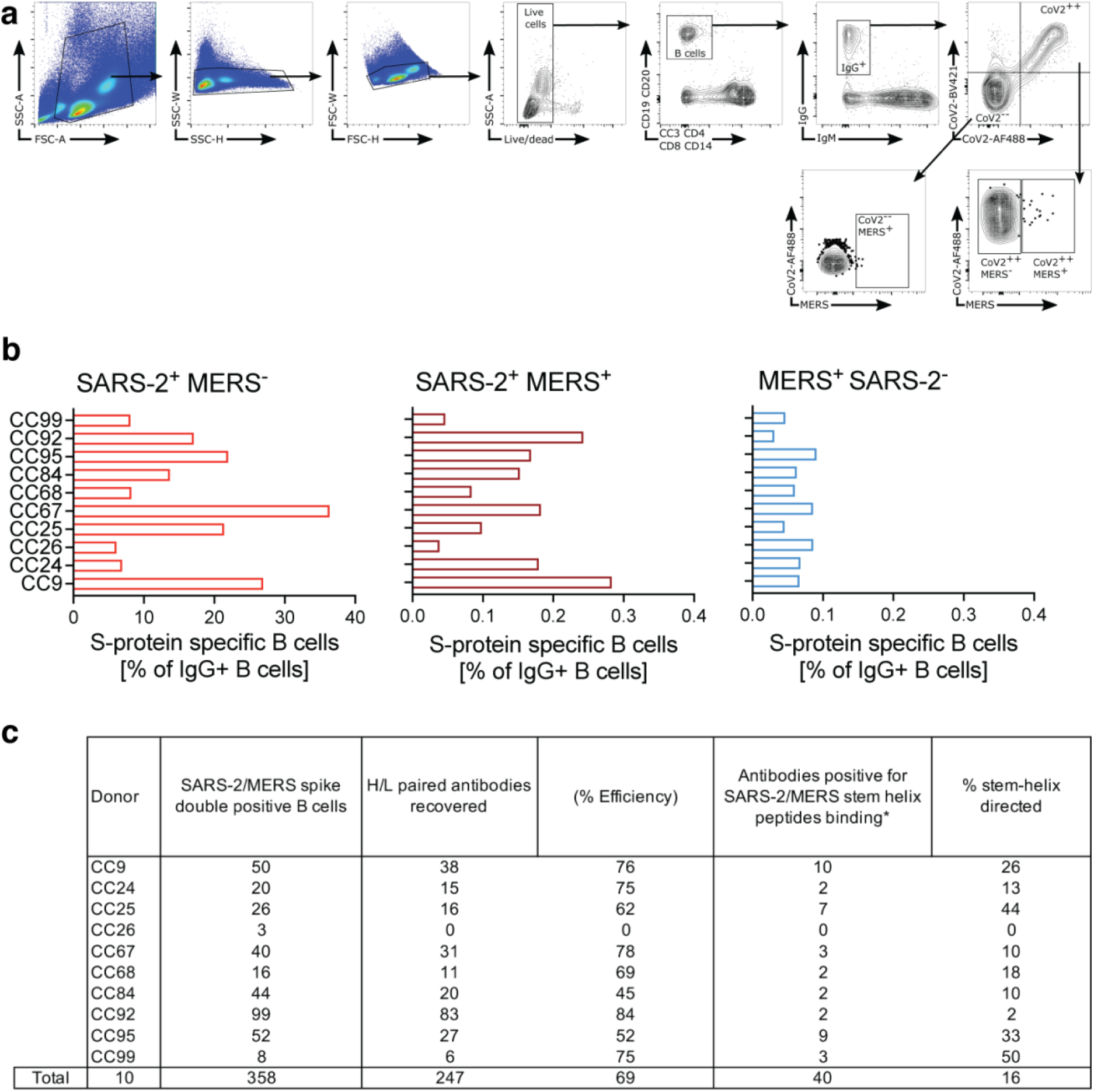
Flow cytometry B cell profiling, sorting strategy and SARS-CoV-2 and MERS-CoV S-protein specific B cells in infected-vaccinated donors. **a.** Gating strategy for analysis of IgG^+^ B cell populations that bind MERS-CoV S-protein only (CD3^−^CD4^−^CD8^−^CD14^−^CD19^+^CD20^+^IgM^−^IgG^+^CoV2^−−^MERS-CoV^+^), SARS-CoV-2 S-protein only (CD3^−^CD4^−^CD8^−^CD14^−^CD19^+^CD20^+^IgM^−^IgG^+^CoV2^++^MERS-CoV^−^), or both MERS-CoV and SARS-CoV-2 S-proteins (CD3^−^CD4^−^CD8^−^CD14^−^CD19^+^CD20^+^IgM^−^IgG^+^CoV2^++^MERS-CoV^+^). **b.** The frequencies of SARS-CoV-2 S-protein-specific IgG^+^ B cells (left), SARS-CoV-2 and MERS-CoV double positive S-protein-specific IgG^+^ cross-reactive B cells (middle) or MERS-CoV S-protein-specific IgG^+^ B cells (right) in PBMCs of 10 infected vaccined-vaccinated donors. **c.** Summary of the number of SARS-CoV-2 and MERS-CoV double positive S-protein specific cross-reactive B cells recovered from each of the donor, number and efficiency of heavy and light chain paired recovered, number of stem-helix mAb in each donor and their frequency out of the S-protein specific cross-reactive IgG B cells.

**Supplementary Figure 2.**
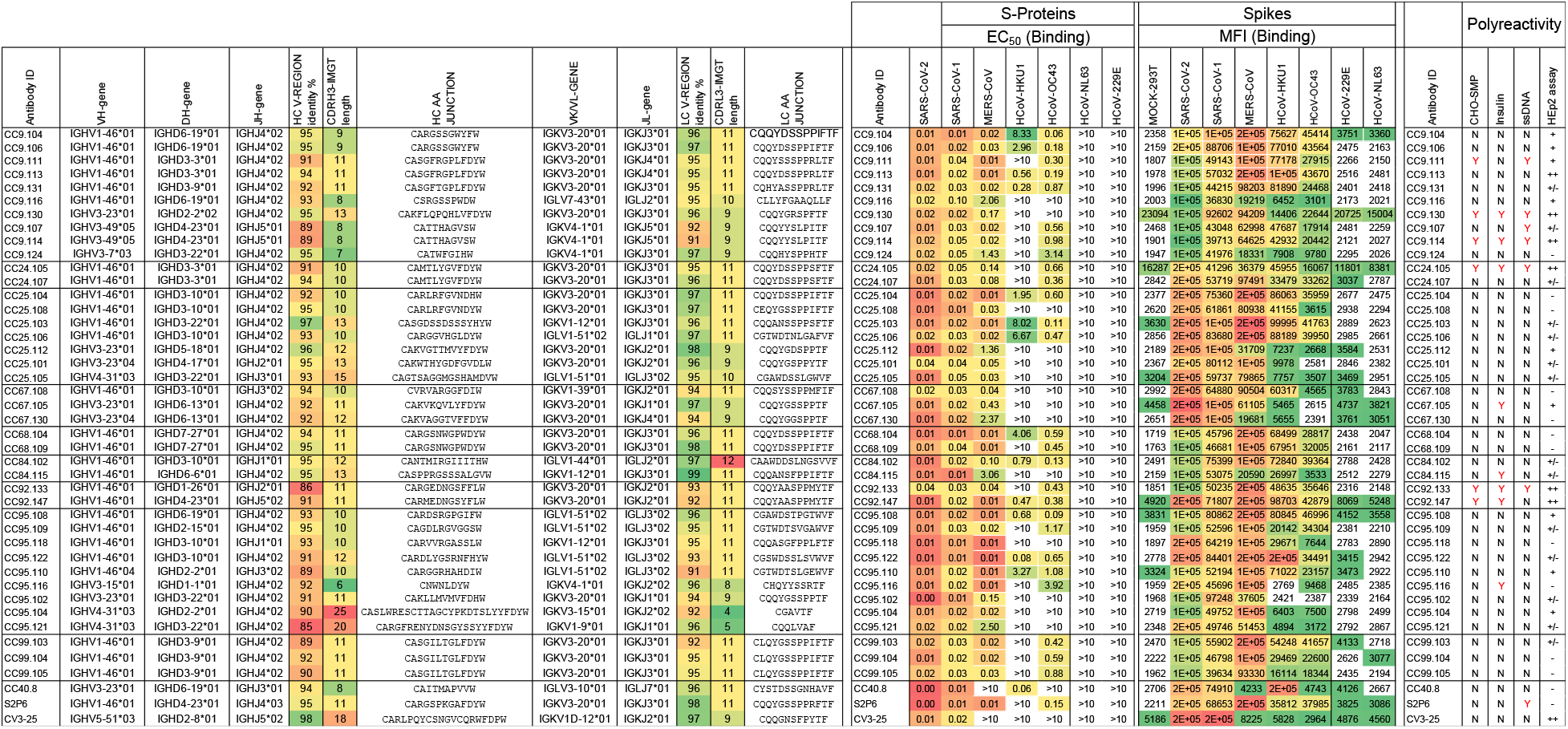
Binding and immunogenetic properties of the isolated S2 stem-helix mAbs. A total of 40 S2 stem-helix mAbs from 9 SARS-CoV-2 infected-vaccinated donors (CC9 (n = 10), CC24 (n = 2), CC25 (n = 7), CC67 (n = 3), CC68 (n = 2), CC84 (n = 2), CC92 (n = 2), CC95 (n = 9) and CC99 (n = 3) were isolated by single B cell sorting using SARS-CoV-2 and MERS-CoV S-proteins as baits. 32 out of 40 mAb were encoded by unique gene families. Heavy (V, D, J) and light (V, J) germline gene usage, CDR3 lengths and somatic hypermutation (SHM) levels are shown. MAbs were expressed and tested for binding to soluble (ELISA) and cell surface expressed (Cell-ELISA) spikes derived from human β-(SARS-CoV-1 or 2, MERS-CoV, HCoV-HKU1 and HCoV-OC43) and α-(HCoV-NL63 and HCoV-229E) coronaviruses and EC_50_ and MFI (mean fluorescent intensity) binding values are shown. S2 stem-helix bnAbs show binding to β-but not α-HCoV spikes. Binding to cell surface expressed spikes was relatively better compared to soluble S-proteins. Polyreactive binding analysis of S2 stem-helix bnAbs to HEp2 cells and by ELISA for binding against polyspecific reagents (PSR) including Chinese hamster ovary cells solubilized membrane protein (CHO-SMP), insulin and single-strand DNA (ssDNA). More details are included in fig. S4. S2 stem helix bnAbs, CC40.8, S2P6 and CV3-25 were used as control for binding assays.

**Supplementary Figure 3.**
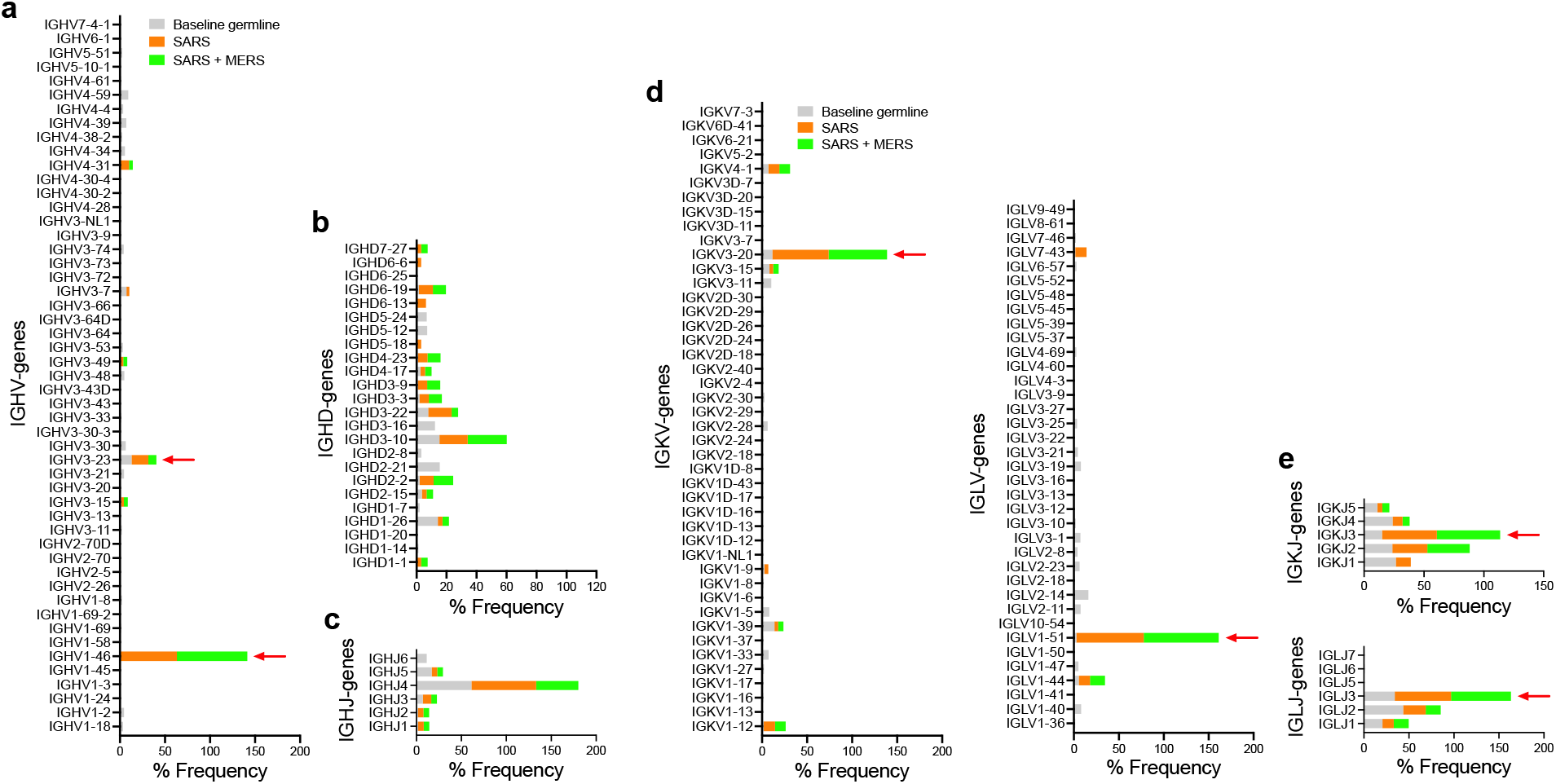
Immunoglobulin heavy and light chain gene usage and enrichment in isolated mAbs compared to a reference human germline database. Baseline germline frequencies of heavy chain genes (IGHV, IGHD and IGHJ genes) **(a., b., c)** and light chain genes (IGKV, IGLV, IGKJ and IGLJ genes) **(d., e)** are shown in grey, and S2 stem helix sarbecovirus bnAbs (SARS: orange) and sarbecovirus + MERS-CoV bnAb (SARS + MERS: green) are shown. Arrows indicate gene enrichments compared to human baseline germline frequencies.

**Supplementary Figure 4.**
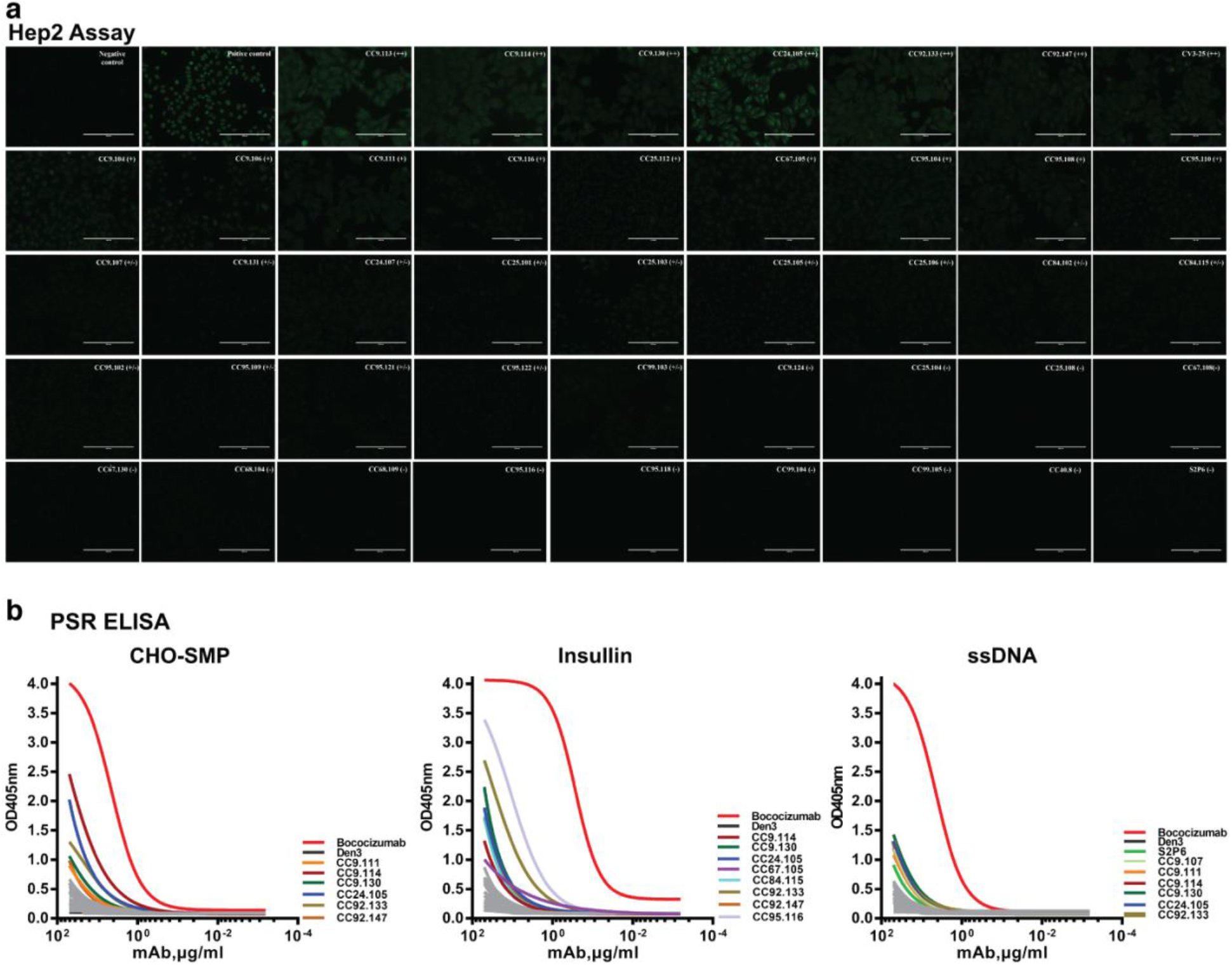
Evaluation of stem-helix bnAbs for polyreactivity and autoreactivity. **a-b.** Antibodies were tested for binding to immobilized HEp2 cells **(a)** and by ELISA for binding against polyspecific reagents (PSR) including Chinese hamster ovary cells solubilized membrane protein (CHO-SMP), insulin and single-strand DNA (ssDNA) **(b)**. For HEp2 assay, immunofluorescence showed binding of antibodies to immobilized HEp2 cells was detected by FITC-labelled secondary antibody. Fluorescent intensity from strong to weak were labeled as “++”, “+” and “+/−” accordingly. “−” indicated little or no signal could be observed. Positive and negative controls for the HEp2 assay are provided by the manufacturer. In PSR ELISA, Bococizumab which is a humanized mAb targeting the LDL receptor-binding domain of PCSK9 and studied in phase I–III clinical studies (*1*), was used as a positive control. The color curves indicate antibodies that can react with PSR, while gray curves are the antibodies with little or no binding to PSR. DEN3 mAb was used as a negative control.

**Supplementary Figure 5.**
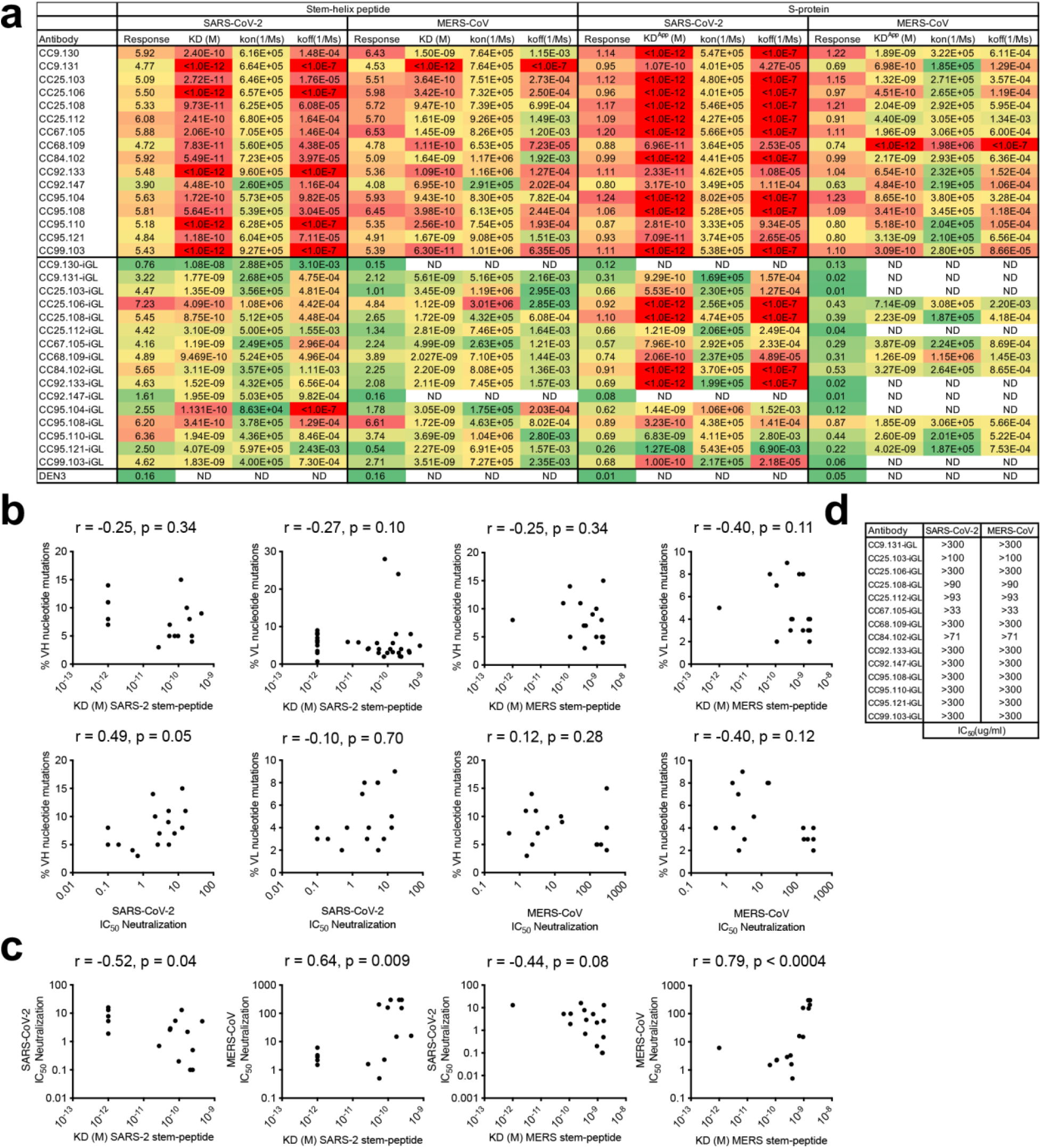
BLI Binding of S2 stem bnAbs and their iGLs with SARS-CoV-2 and MERS-CoV stem-helix peptides and S-proteins and association with SHMs and Neutralization. **a.** BioLayer Interferometry (BLI) binding kinetics of 16 S2 stem-helix bnAbs and their inferred germline (iGL) Ab versions with SARS-CoV-2 and MERS-CoV stem-helix peptides and S-proteins. Binding kinetics were obtained using the 1:1 binding kinetics fitting model on ForteBio Data Analysis software and maximum binding responses, dissociations constants (*K*_D_) and on-rate (*k_on_*) and off-rate constants (*k_off_*) for each antibody peptide interaction are shown. K_D_, k_on_ and k_off_ values were calculated only for antibody-antigen interactions where a maximum binding response of 0.2nm was obtained. MAbs were also tested with SARS-CoV-2 and MERS-CoV S-proteins and the responses, apparent binding constants (K_D_^App^) and *k_on_* and *k_off_* constants for each antibody-antigen interaction are indicated. The iGL Ab versions of stem-helix bnAbs showed reduced binding compared their mature versions. **b.** Correlations of stem-helix mAb binding (*K*_D_ (M) values) to SARS-CoV-2 and MERS-CoV peptides and virus neutralization with heavy (VH) chain and light (VL) chain SHM levels. **c.** Correlations of stem-helix mAb binding (*K*_D_ (M) values) to SARS-CoV-2 and MERS-CoV peptides with neutralization against their corresponding viruses. Correlations were determined by nonparametric Spearman correlation two-tailed test with 95% confidence interval. The Spearman correlation coefficient (r) and p-value are indicated. **d.** IC_50_ neutralization of S2 stem-helix bnAb iGLs with SARS-CoV-2 and MERS-CoV.

**Supplementary Figure 6.**
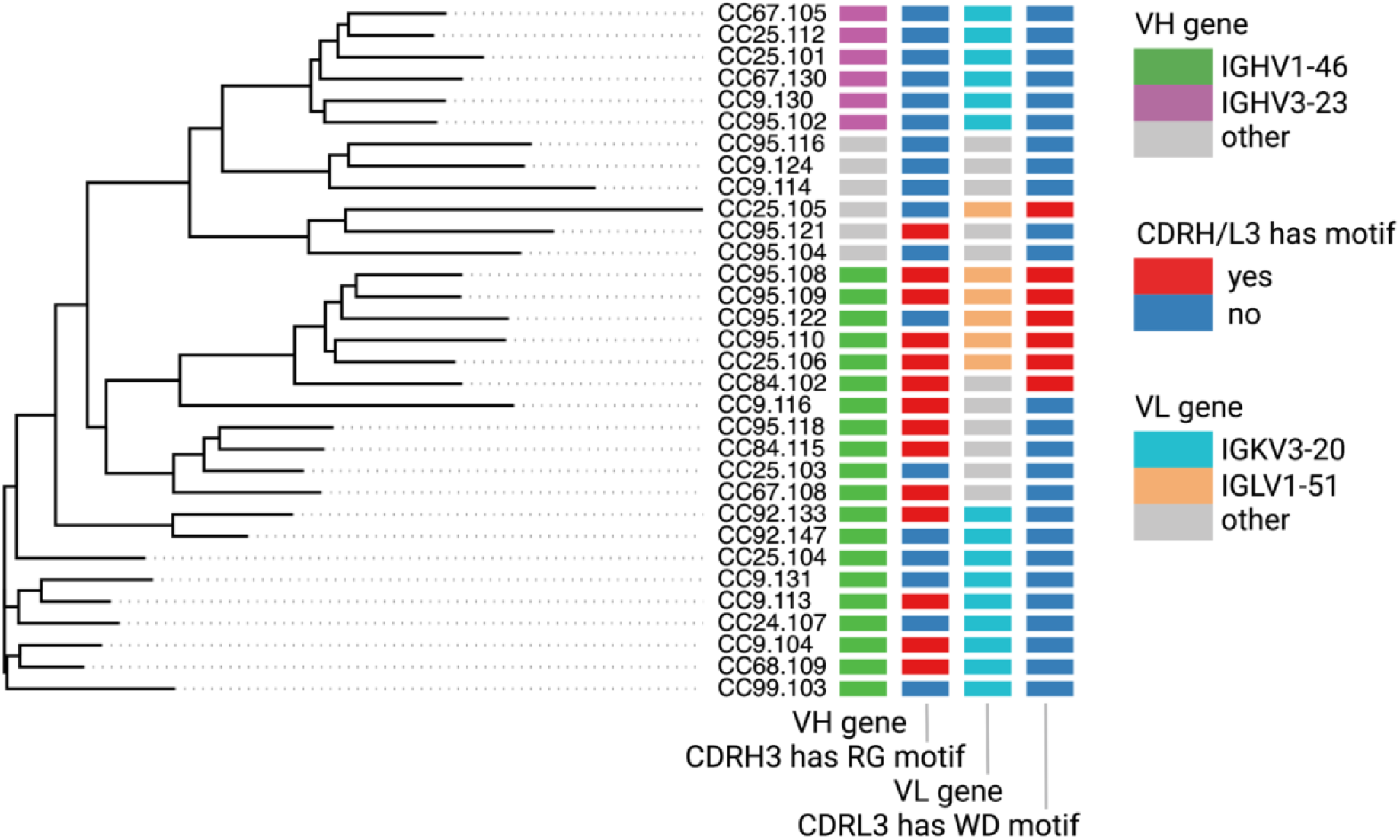
Immunogenetics analysis of heavy and light chain sequences of 32 unique S2 stem-helix mAbs. The phylogenetic tree represents concatenated heavy and light chain amino acid sequences of 32 S2 stem-helix mAbs. mAbs IDs are shown on the right. Four colored columns on the right show the following characteristics of mAbs (from left to right): (1) the germline V gene of each heavy chain (IGHV1-46: green, IGHV3-23: plum, others: gray), (2) the presence of RG motif in the amino acid sequence of each CDRH3 (motif is present: red, motif is missing: blue), (3) the germline V gene of each light chain (IGKV3-20: sky, IGLV1-51: cantaloupe, others: gray), (4) the presence of WD motif in the amino acid sequence of each CDRL3 (motif is present: red, motif is missing: blue).

**Supplementary Figure 7.**
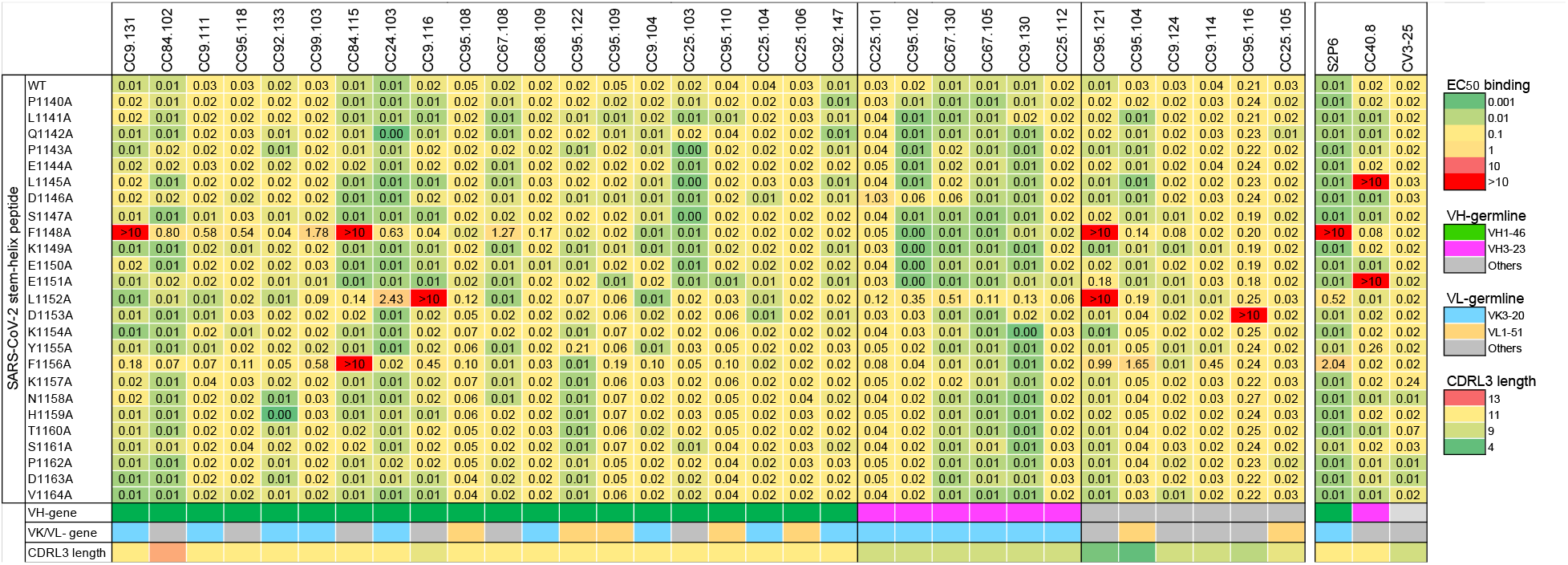
Epitope mapping of S2 stem-helix bnAbs with SARS-CoV-2 stem-helix peptide alanine scan mutants. Heatmap showing EC__50__ ELISA binding titers of S2-stem helix bnAbs to 25mer SARS-CoV-2 stem-helix peptide and its alanine scan mutants. Three hydrophobic residues, F^1148^, L^1152^ and F^1156^ were commonly targeted by stem-helix bnAbs. S2 stem-helix bnAbs are grouped based on their heavy chain gene usage (IGHV1-46, IGHV3-23 and others). The light chain germline genes (IGKV3-20, IGLV1-51 and other) and CDRL3 lengths are shown. S2P6, CC40.8 and CV3-25 S2 stem-helix mAbs were used as controls.

**Supplementary Figure 8.**
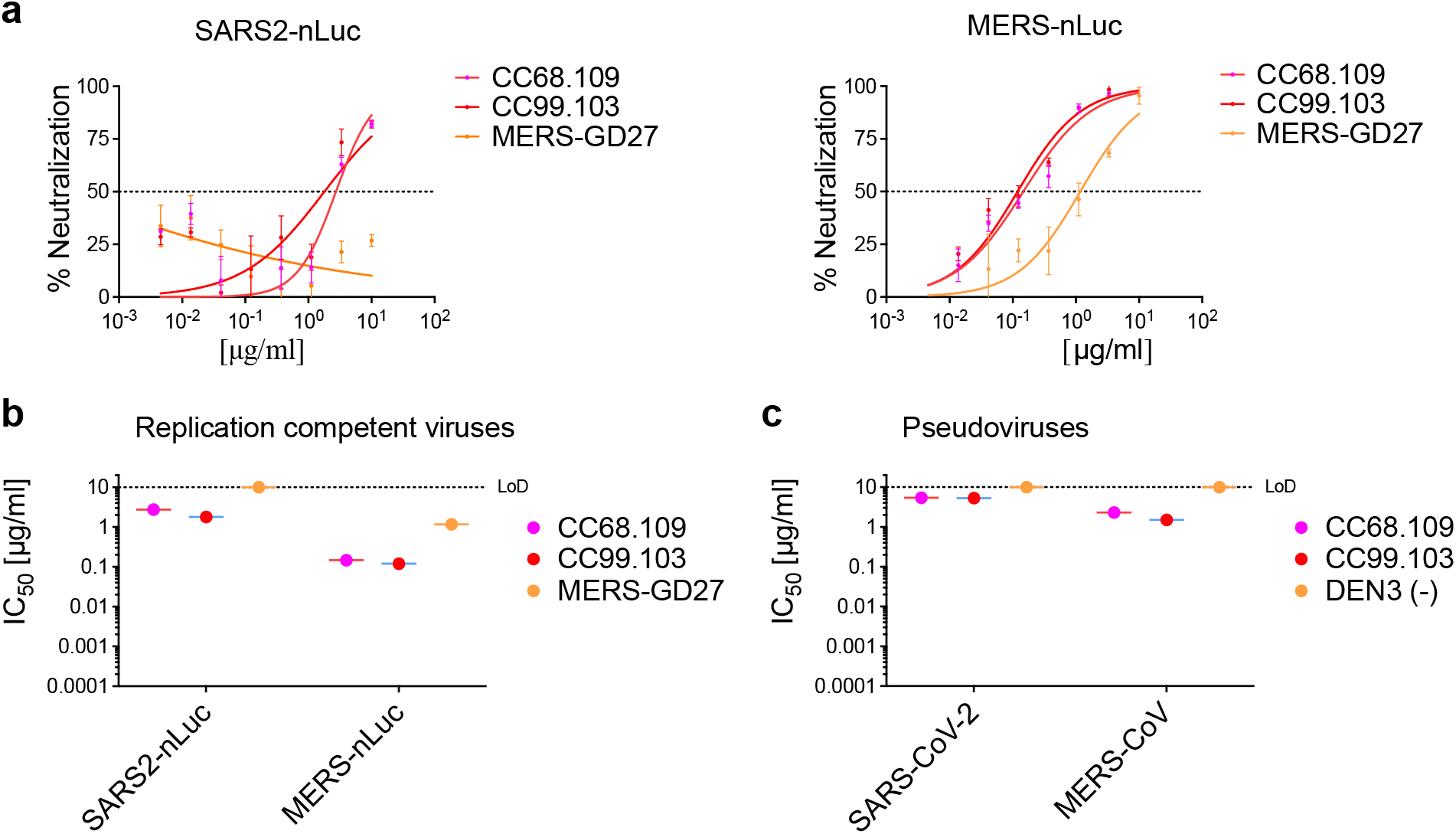
Neutralization of replication competent betacoronaviruses by select S2-stem helix bnAbs. **a.** Neutralization of replication competent viruses encoding SARS-CoV-2 (SARS2-nLuc), and MERS-CoV (MERS-nLuc) by 2 select S2 stem-helix bnAbs, CC68.109, and CC99.103. MERS-GD27 antibody (*2*) was a positive control for the MERS-CoV neutralization assay. **b-c.** Comparison of IC_50_ neutralization titers of S2 stem-helix bnAbs with replication-competent **(b)** and pseudoviruses **(c)** of SARS-CoV-2 and MERS-CoV.

